# Cross-*strata* co-occurrence of ripples with theta-frequency oscillations in the hippocampus of foraging rats

**DOI:** 10.1101/2022.11.15.516579

**Authors:** Pavithraa Seenivasan, Reshma Basak, Rishikesh Narayanan

## Abstract

**Background and motivation:** Brain rhythms have been postulated to play central roles in animal cognition. A prominently reported dichotomy of hippocampal rhythms, driven primarily by historic single-*strata* recordings, assigns theta-frequency oscillations (4–12 Hz) and ripples (120–250 Hz) to be exclusively associated with preparatory and consummatory behaviors, respectively. However, due to the differential power expression of these two signals across hippocampal *strata*, reports of such exclusivity require validation through simultaneous multi-*strata* recordings and cross-*strata* analysis of these oscillatory patterns.

**Methodology:** We assessed co-occurrence of theta-frequency oscillations with ripples in multi-channel recordings of extracellular potentials across hippocampal *strata* from foraging rats. We detected all ripple events from an identified *stratum pyramidale* (*SP*) channel based on rigorous thresholds relating to the spectro-temporal and spatial characteristics of ripples. We then defined theta epochs based on theta oscillations detected from each of the different channels spanning the *SP* to the *stratum lacunosum-moleculare* (*SLM*) through the *stratum radiatum* (*SR*). We calculated the proportion of ripples embedded within theta epochs.

**Results:** We found ∼20% (across rats) of ripple events (in *SP*) to *co-occur* with theta epochs identified from *SR*/*SLM* channels, defined here as *theta ripples*. All characteristics of theta ripples were comparable with ripples that occurred in the absence of theta oscillations. Furthermore, the power of theta oscillations in the immediate vicinity of theta ripples was similar to theta power across identified theta epochs, together validating the identification process of theta ripples. Strikingly, when theta epochs were instead identified from the *SP* channel, such co-occurrences were significantly lower in number. The reduction in the number of theta ripples was consequent to progressive reduction in theta power along the *SLM-SR-SP* axis. We assessed the behavioral state of rats during ripple events and found most theta ripples to occur during immobile periods. We confirmed that across sessions and rats, the theta power observed during exploratory theta epochs was comparable with theta power during immobile theta epochs. In addition, the progressive reduction in theta power along the *SLM-SR-SP* axis was common to both exploratory and immobile periods. Finally, we found a strong theta-phase preference of theta ripples within the third quadrant [3π/2–2π] of the associated theta oscillation.

**Implications:** Our analyses provide direct quantitative evidence for the occurrence of ripple events nested within theta oscillations in the rodent hippocampus. These analyses emphasize that the prevalent dichotomy about the manifestation of theta-frequency oscillations and ripples needs to be reevaluated, after explicitly accounting for the differential *stratum*-dependent expression of these two oscillatory patterns. The prevalence of theta ripples expands the potential roles of ripple-frequency oscillations to span the continuum of encoding, retrieval, and consolidation, achieved through interactions with theta oscillations.

## INTRODUCTION

Historically guided by single *strata* recordings, a strong narrative has driven a dichotomy in the expression of theta-frequency oscillations and ripples in the rodent hippocampus. Specifically, from a behavioral perspective, theta oscillations (4–10 Hz) are associated with exploration or active engagement with external stimuli. On the contrary, ripple events (120– 250 Hz) embedded within sharp wave ripple (SPW-R) complexes have been observed during slow wave sleep (SWS) or wakeful immobility. In addition to this behavioral dichotomy, theta oscillations are implicated in encoding phenomena whereas ripples are associated with memory consolidation and retrieval. The exclusive nature of this dichotomy, in terms of the two distinct oscillatory bands being involved in behavioral and physiological processes, provides a compelling narrative for how electrical signals drive memory processes (Buzsaki, 1989, 2002; Klausberger et al., 2003; Girardeau and Zugaro, 2011; Colgin, 2013; Hasselmo and Stern, 2014; Buzsaki, 2015; Roumis and Frank, 2015; Joo and Frank, 2018; Girardeau and Lopes-Dos-Santos, 2021). In this study, we systematically assessed this dichotomy using simultaneous multi-*strata* recordings from hippocampal CA1 of foraging rats, specifically asking if the occurrence of theta-frequency oscillations and ripples are indeed mutually exclusive in the recorded local field potentials (LFP).

Our rationale for revisiting the dichotomy and probing co-occurrences of theta- and ripple-frequency oscillations is three-fold. First, there are strong lines of evidence in the literature that the power distribution of theta-frequency oscillations and ripples are differential across different *strata* of the CA1 subregion (Buzsaki, 2002; Gordon et al., 2005; Buzsáki, 2015; Navas-Olive et al., 2022; Zutshi et al., 2022). Whereas theta-frequency oscillations are prominent in the *stratum lacunosum-moleculare* (SLM) and the distal *stratum radiatum* (SR), ripples are rich within the *stratum pyramidale* (SP). From an experimental design perspective, the differential power distribution of these two signals (theta-frequency oscillations and ripples) implies that their co-occurrence should not be assessed with single *strata* recordings, but with simultaneous multi-*strata* recordings that span the SLM, SR and SP. From an analysis standpoint, this differential distribution requires that the two signals are detected from the *strata* where their *respective* powers are higher. In other words, to analyze co-occurrence of these two oscillatory patterns, it is essential that ripples are detected from the SP LFP and theta oscillations are identified from the SLM/SR LFP. The differential power distribution implies that detection of ripples from SLM/SR or theta oscillations from the SP would result in lower numbers of both identified oscillatory patterns, thus providing a false dichotomy for their exclusive expression. Therefore, we hypothesized that several co-occurrences of theta oscillations with ripples in simultaneous multi-*strata* recordings would manifest if each oscillatory band were identified from the *strata* where they express with the highest power. Testing this hypothesis necessitated simultaneous acquisition of extracellular signals across multiple *strata* coupled with an analysis pipeline that identifies theta oscillations and ripples from the *stratum* where their power is known to be high.

Second, there are some pointers in the literature that suggest the possibility of ripples co-occurring with theta rhythm during REM sleep (Fig. 5 in (Buzsáki, 2015)) as well as during exploratory periods (O’Neill et al., 2006). However, these studies did not investigate ripples from the perspective of their cross-*strata* temporal coexistence with theta oscillations. Finally, there are lines of evidence for a shared role for plateau potentials (consequent to strong dendritic depolarization) during theta oscillations as well as in driving sharp waves in the CA1 subregion (Buzsáki and Vanderwolf, 1983; Kamondi et al., 1998a; Epsztein et al., 2011; Bittner et al., 2015; Buzsaki, 2015; Bittner et al., 2017; Zhao et al., 2020; McKenzie et al., 2021; Milstein et al., 2021). This shared cellular signature provided us a further motivation to assess the potential co-occurrence of ripples with theta oscillations.

Motivated by these observations, we systematically assessed the hypothesis that theta oscillations and ripples might co-occur across hippocampal CA1 *strata*, given the established *stratum-*dependent expression profiles of LFPs in these two oscillatory bands. In testing the hypothesis, we introduced important changes to the experimental design and analyses pipeline to explicitly account for these differential expression profiles. First, we simultaneously recorded extracellular signals from multiple *strata* using silicon polytrodes in awake behaving animals, rather than restricting our attention to one specific *strata*. Second, we *independently* analyzed electrophysiological and behavioral data instead of making strong assumptions about the occurrence of specific bands of oscillations during specific behavior. In doing this, we placed tight constraints (with reference to frequency and durations) on the identification of ripples and theta epochs, as well as on the delineation of immobile *vs*. exploratory periods of behavior (with reference to velocity and durations). Third, and most importantly, we chose *different strata* for analyzing theta oscillations and ripples, explicitly based on established knowledge about where the powers of these two oscillatory bands are maximal. Specifically, we chose the SP for identifying ripples and the SLM/SR for delineating theta epochs. We then assessed the cross-*strata* co-occurrence of theta oscillations and ripples using these simultaneously recorded LFP waveforms.

Our experimental design and analysis pipeline that accounted for differential expression profiles unveiled the existence of *theta ripples*, defined as ripples in the SP that co-occurred with theta oscillations in the SLM/SR. Around 20% of all detected ripples were theta ripples and occurred predominantly during immobile periods of foraging rats. All characteristics of theta ripples were comparable with those ripples that occurred in the absence of theta oscillations. Furthermore, the power of theta oscillations in the immediate vicinity of theta ripples was similar to theta power across identified theta epochs, together validating the identification process of theta ripples. Importantly, our analyses showed that theta oscillations observed during immobile and exploratory periods were comparable in their power across *strata*, with a *stratum*-dependent reduction in the theta along the SLM-SR-SP-SO (SO stands for *stratum oriens*) axis for both periods. As a final test of our hypothesis, we used different channels along the SLM-SR-SP-SO axis to identify theta epochs to count the number of SP ripples counted as theta ripples when each of these channels were used as theta channels. We found a significant reduction in the number of identified theta ripples when SP or SO channel was chosen as the theta channel, compared to when SLM was the theta channel.

Our study demonstrates that the differential power profile stipulates simultaneous multi-*strata* recordings and advocates for the use of different *strata* for identifying theta oscillations and ripples, explicitly based on where their power is maximal. Deviations from the *strata* where these signals of two different frequencies have their highest power do not account for the differential spatial expression and result in missing co-occurrences of these two oscillations. Together, we argue that the prevalent dichotomy about the manifestation of theta-frequency oscillations and ripples should be reevaluated, after explicitly accounting for the differential *stratum*-dependent expression of these two oscillatory patterns. The prevalence of theta ripples expands the potential roles of ripple-frequency oscillations to span the continuum of encoding, retrieval, and consolidation, achieved through interactions with theta oscillations.

## MATERIALS AND METHODS

### Surgical procedures

Male Sprague Dawley (SD) rats of 6–8 months age were entrained to a 14-hour light, 10-hour dark cycle for at least two weeks prior to surgery and were housed one per cage before and after surgery. They were provided *ad libitum* food and water, with the housing temperature maintained between 21–23° C. All experimental procedures were approved by the Institutional Animal Ethics Committee of the Indian Institute of Science, Bangalore.

Animals were anesthetized with gaseous isoflurane throughout the course of surgery and craniotomies were performed with a stereotaxic apparatus (David Kopf Instruments, 963LS). The coordinates for craniotomy were fixed at 3.5 mm along the anterior-posterior (AP) axis from the bregma and 2.2 mm medio-laterally (ML) from the central axis. Silicon probes, specifically, polytrodes, with 32 contact points and 25 μm spacing between the contact points (Neuronexus Inc., A1x32-Poly2) were lowered ∼3 mm dorso-ventrally (DV) from the surface of the brain, following durectomy to reach the dorsal hippocampal CA1 (Fig. S1). The lowering process was done at the rate of ∼1 μm/s to minimize probe and tissue damage as well as to facilitate smooth implantation. The craniotomy was then sealed with sterile bone wax. Six screws were drilled through the skull prior to craniotomy, to stabilize and strengthen the implant, two of which in the cerebellum served as ground and reference electrodes. The exposed skull area was then built up with dental cement all the way from the skull surface up to the base of the probe to provide stability and long-term retention of the implant.

### Electrophysiological and behavioral acquisition

The rats were allowed to recover for about a week following surgery, during which they were gentled and habituated to the experimenter, recording room and acquisition cables. Recordings began the following week, during which extracellular field potentials were recorded from the hippocampus while the rats were engaged in a foraging behavior. These extracellular potentials were acquired using a Neuralynx acquisition system at a sampling rate of 32 kHz and with a band pass frequency range of 0.1–6 kHz.

Behavior involved exploration and foraging in an open cardboard box of dimensions 59 cm × 49 cm × 47 cm (Fig. S1). The box was lined with *clingwrapp* plastic on the inside and a copper mesh on the outside (Fig. S1). The interior plastic lining facilitated cleaning the arena walls and minimized the retention of olfactory cues. The exterior copper mesh which was grounded to a thick copper plate served as a Faraday cage to reduce external electromagnetic interference. Rats foraged in 3 kinds of arenas, distinguished from each other by three differently textured mats fixed as the floor of the arena. On a given day, animals foraged in 2 different arenas with 3 consecutive sessions in each arena, resulting in a total of 6 sessions per day. Each recording session lasted for a duration of 15–35 minutes and a total of 6 days of recordings were performed for each rat, thus giving a total of 36 sessions per rat. Four pieces of sweet biscuits (threptin diskettes, ∼0.1 g weight) were manually pre-placed in random locations within the arena by the experimenter. The recording box that was lined with plastic on the inside was ethanol-wiped and cleaned between arena switches (once every 3 sessions), while the arena mats were washed following a consecutive 3-session usage. Recordings were consistently performed around the same time on any day, during their active time in complete darkness. Behavioral recordings were also performed in a linear maze set up for 2 rats (Rats 2 & 3). This consisted of a rectangular, laminated wooden box with dimensions, 98 cm × 17 cm × 24 cm. Each rat had 2 days of recordings in the linear maze, with 3 sessions (15–20 minutes per session) per day separated by a 5-minute break between sessions.

Behavioral data was captured in the dark as video frames at a sampling rate of 60 frames per second, using a camera (Sony PlayStation 3 eye, 99028) that was positioned on top of the arena. Bonsai (Lopes et al., 2015) was used to set up the behavioral recording pipeline and served as an interface between the electrophysiological and behavioral acquisition. We employed custom-written programs in Bonsai to identify the LED headlight attached to the Neuralynx headstage, against a dark background based on color thresholding. This allowed us to track the centroid of the LED as a function of time and to obtain the trajectory of the rat through each session of recording. We employed Bonsai to communicate with the camera as an input device connected to the computer while it connected with Neuralynx through a digital output signal that was sent through Arduino. A record of the timing of this digital output signal from the camera was saved along with the electrophysiological data in Neuralynx, along with the positional information from the camera. This common origin time aided in referencing the two clocks towards synchronizing electrophysiology with behavior during offline analyses.

### Preprocessing of raw signals

All analyses of the acquired extracellular field recordings were performed using custom-written script within the FMA toolbox (https://fmatoolbox.sourceforge.net/). Raw signals, sampled at 32 kHz, were decimated to 4 kHz and filtered between 0.5–300 Hz using a zero phase Butterworth filter of order 4, to derive local field potentials (LFP). Movement and grooming artifacts in the signal were identified as fluctuations that exceeded 5 standard deviations of the baseline signal and were removed.

### Identification of hippocampal *strata*

The various hippocampal CA1 *strata* – *stratum oriens* (SO), *stratum pyramidale* (SP), *stratum radiatum* (SR), and *stratum lacunosum-moleculare* (SLM) – were identified using multiple strategies. First, we used histological verification to infer the approximate location of the 800 μm silicon probe in the hippocampus of all rats (Fig. S2*A*). Second, we assessed the laminar profiles of LFP (0.5–300 Hz) signals to identify *strata* using well-established *strata*-specific characteristic signatures (Fig S2*B*). To elaborate, sharp waves and ripples that are part of sharp wave ripple (SPW-R) complexes are known to spatially localize themselves within distinct CA1 *strata*. Specifically, ripples express themselves within the SP layer whereas sharp waves are observed in the SR layer of the CA1 (Gordon et al., 2005; Mizuseki et al., 2011; Buzsáki, 2015; Navas-Olive et al., 2022). We therefore marked the channel with maximal number of ripples as the SP and that with prominent sharp waves as the SR layer. Additionally, the SR– SLM band was identified by virtue of enhanced power in the theta frequency (6–10 Hz) band, the ordered organization of CA1 *strata* as SO-SP-SR-SLM (Andersen et al., 2006), and the fixed distances between recording channels in our electrode (25 μm). SO was marked to include channels 100–150 μm above SP. Third, we confirmed *strata* identification through phase shifts observed in theta oscillations across different channels (Fig S3). In accordance with the characteristic phase inversion that occurs from SO to SLM, we noted that the phase of the recorded signals reversed close to the identified SP layer across all 3 rats (Fig. S3).

### Detection of theta epochs and ripple events

Theta epochs were detected using instantaneous power within the theta frequency band (default 6–10 Hz) computed through multi-taper spectral estimation (Thomson, 1982). They were identified as segments of the recorded voltage signal where power in the theta frequency band exceeded 1 SD above mean. To enforce strictness in the identification of theta epochs, the minimum duration necessary for detection as a theta epoch was fixed to be 1 second. Analyses with different procedures for identifying theta epochs were employed to enable further corroboration of our results (Table S1–S3).

To detect ripple events, the LFP signals recorded from all the channels were band-pass filtered in the ripple frequency range (120–250 Hz) using a zero-phase Butterworth filter of order 4. This filtered signal was subjected to ripple identification criteria. Specifically, the beginning and the end of a ripple were detected based on instances of ripple power exceeding 2 SD above mean and the peak of the ripple was identified by a 5 SD deflection above mean. The minimum and maximum durations of ripple events were fixed as 20 and 200 ms respectively, and the minimum inter-ripple interval was fixed at 30 ms. The entire ripple identification procedure was independently applied to all the channels initially, to identify *strata* and to eliminate events detected across many channels that are likely to be artifacts. Specifically, after detecting ripple-like events in all the recorded channels, putative ripple events in the SP layer were discarded if there were events detected in at least 10 channels within 10 ms on either side of the ripple under consideration (*e*.*g*., Fig. S4). These events, owing to their simultaneous detection across channels, are likely to be related to movement or muscle related artifacts (*e*.*g*., Fig. S4) and not real ripples. For all analyses reported in this study, we considered ripple events detected in the channel designated as *stratum pyramidale*.

### Defining, classifying, quantifying, and validating different types of ripples

Theta ripple (θ-ripple) co-occurrences were defined as events wherein a ripple detected in SP was temporally localized within a theta epoch identified from recordings from the SR or SLM layers. Such a cross channel approach was used for the detection of theta ripples, owing to their differential expression across hippocampal *strata* (Gordon et al., 2005; Mizuseki et al., 2011; Buzsáki, 2015; Navas-Olive et al., 2022), with ripples and theta oscillations predominantly observed in SP and SR/SLM respectively. This results in a physiological characterization of ripples based on whether they belonged to a theta epoch or not. These theta epochs that were identified independently were extended to 100 ms on either side, so that instances wherein ripple peak times were just outside the theta epochs while the start of the ripple still fell within the theta epoch were not neglected.

#### Cross-validating theta oscillations and ripples

Ripples that occurred along with theta oscillations were validated by quantifying several signature characteristics of ripples and comparing them with the classic non-theta ripples. The amplitude of ripples was computed as the difference between the maximal and minimal voltage values within each ripple waveform. The duration of ripples was computed as the difference between the detected start and end timestamps of a ripple. Power spectral density (PSD) analysis involved computing the Fourier transform profiles of both theta and non-theta ripple waveforms. The frequency at which the ripple PSD manifested maximal power was computed for every considered ripple waveform and averaged across all theta ripples and non-theta ripples independently. All other ripple measurements were performed on all theta and non-theta ripples. For PSD analysis, 10% of ripples from each session were chosen in a pseudorandom fashion to assess the spectral characteristics of theta *vs*. non-theta ripples.

Theta oscillations observed as part of theta ripples were validated by comparing their spectral power with theta power during classic theta epochs. Theta power was computed as the power in the theta frequency band and was computed for each detected theta epoch.

#### Behavioral categorization of ripples

Positional information of the rat was continuously available from time-synced video recordings of foraging behavior. The instantaneous linear velocity of the rat was computed based on the Euclidean distance between successive positional coordinates tracked by the camera. This instantaneous velocity was used to characterize two kinds of behavioral epochs during foraging: exploratory epochs were periods with a detected running velocity > 5 cm/s and immobile periods were those where the running velocity < 5 cm/s. To ensure strict classification of ripples to behavioral states, immobile or exploratory periods were required to exceed a minimum duration of 1 s to be assigned to the respective epochs. As our classification of behavioral epochs were strictly based on velocity and duration, there were ripples that were not classified into either immobile or exploratory epochs. This behavioral categorization (exploratory *vs*. immobile behavioral epochs), in addition to the electrophysiological characterization (theta *vs*. non-theta ripples) resulted in four different kinds of ripples that are based on electrophysiology and behavior: exploratory theta (exp. θ); exploratory non-theta (exp. non-θ); immobile theta (imm. θ); and immobile non-theta (imm. non-θ).

#### Fraction and frequency of ripples

The proportions of different kinds of ripples in a session were computed as the ratio of the number of ripples belonging to a specific category to the total number of ripples in that session. For example, the fraction of exploratory θ ripples was computed as the ratio between number of exploratory θ ripples and the total number of ripples. The frequency of ripple occurrence was computed as the ratio of the number of ripples in a certain category to the total duration spent by the rat in that category. For instance, the frequency of occurrence of exploratory θ ripples was computed as the number of exploratory θ ripples divided by the total duration of exploratory theta epochs. These measurements were computed for each session and plotted to emphasize the heterogeneities across sessions.

#### Analysis of theta power during ripple periods

To quantify the strength of theta oscillations during ripples, power in the theta frequency band was computed during a 50-ms period on either side of the peak of the detected ripple (total 100 ms). We computed power in the theta frequency band of the SR/SLM channel signals around every detected SP ripple. This theta power was computed as the magnitude of the wavelet spectrum in the 6–10 Hz band that was averaged for the time period. This process was executed for every ripple in every session.

#### Phase analyses

To quantify the temporal relationship between theta oscillations and ripples during co-occurrence events, we first computed the troughs of the theta signal as the local minima of the oscillatory waveform. Then, we computed the phase of the peak of the ripple detected in the SP with respect to the theta oscillation detected in SR/SLM using:

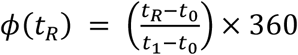

where *t*_*R*_ corresponds to ripple peak time, *t*_0_ and *t*_1_ correspond to theta trough time points that precede and succeed *t*_*R*_, respectively. Specifically, the difference between the ripple peak time and the time of the trough of the theta wave that preceded the ripple peak was normalized by the difference between the times of troughs of the theta wave that encompassed this ripple. This fraction was then multiplied by 360 to compute the phase of the ripple peak time relative to the extracellular theta wave in degrees. We employed this procedure to compute the phases of ripple peak times detected in SP with respect to the theta oscillation from SR/SLM for all theta ripples.

#### Strata-dependence of co-occurrences

We computed the number of co-occurrences of ripples and theta oscillations as a function of the identity of channel that was used to identify theta epochs. Specifically, whereas ripple frequency information was consistently obtained from the identified SP channel, theta epochs were identified independently from all recording channels. The number of theta ripples within each session was computed by calculating the number of ripples that were within theta epochs identified from different recording channels.

#### Long vs. Short ripples

Ripples with a duration > 80 ms were referred to as long ripples while those < 80 ms were considered as short ripples. The fractions of long and short ripples within theta epochs and the fractions of theta and non-theta ripples that were long were computed for each session for all rats.

### Lesion studies and histological verification

To verify the location of surgical implants, we lesioned the region of the implant under anesthesia (ketamine-xylazine) after the completion of all recordings. This was executed by applying a potential difference of 9 V between all 32 electrode sites and a ground site (pinna of the ear) for ∼ 9 s per electrode site. We then performed transcardial perfusion of paraformaldehyde (PFA) following the onset of anesthesia (ketamine-xylazine), after waiting for a day for gliosis to set in. This ensured high contrast staining of the lesioned electrode implant site. The extracted brain was preserved in PFA for 1–2 days following which we prepared coronal slices of 60–150 μm thickness of the implanted hippocampus using a vibratome (Leica). The slices were allowed to dry overnight following which we performed cresyl violet staining. We then mounted slides with DPX mounting medium, left them to dry overnight, and imaged the lesioned implant site of the hippocampus on the following day.

### Details of data and statistical analyses

All data analyses were performed with custom-written scripts that recruited the toolbox (https://fmatoolbox.sourceforge.net/) in Matlab. All statistical analyses were performed using the R computing and statistical software (R Core Team, 2013). To avoid false interpretations and to emphasize the heterogeneities across ripples, sessions, and rats, the entire range of measurements are reported in figures rather than providing only the summary statistics (Marder and Taylor, 2011; Rathour and Narayanan, 2019). The results of statistical tests, with exact *p* values, and the name of the statistical test employed are provided in the figure panels and/or in the respective figure legends.

## RESULTS

### Cross-*strata* co-occurrence of theta frequency oscillations with ripples

Theta-frequency oscillations and ripples are predominantly found in different hippocampal strata. Whereas theta frequency oscillations are rich within the *stratum lacunosum-moleculare* (SLM) and *stratum radiatum* (SR), ripples are prominent in the *stratum pyramidale* (SP) (Gordon et al., 2005). To account for such differential prevalence of different bands of extracellular signals, we simultaneously recorded extracellular field potentials from different hippocampal *strata* of awake behaving rodents that were involved in a foraging task within an open arena (Fig. 1*A*; Fig. S1–S2). Simultaneous acquisition of signals spanning different *strata* using 32-channel silicon polytrodes allowed us to identify each *stratum* using respective characteristic electrophysiological signatures (Fig. S3), and to assess the co-occurrence of valid ripples in the SP (Fig. S4–S8) with theta-frequency oscillations in the SR/SLM (Fig. 1*B*).

**Figure 1.**
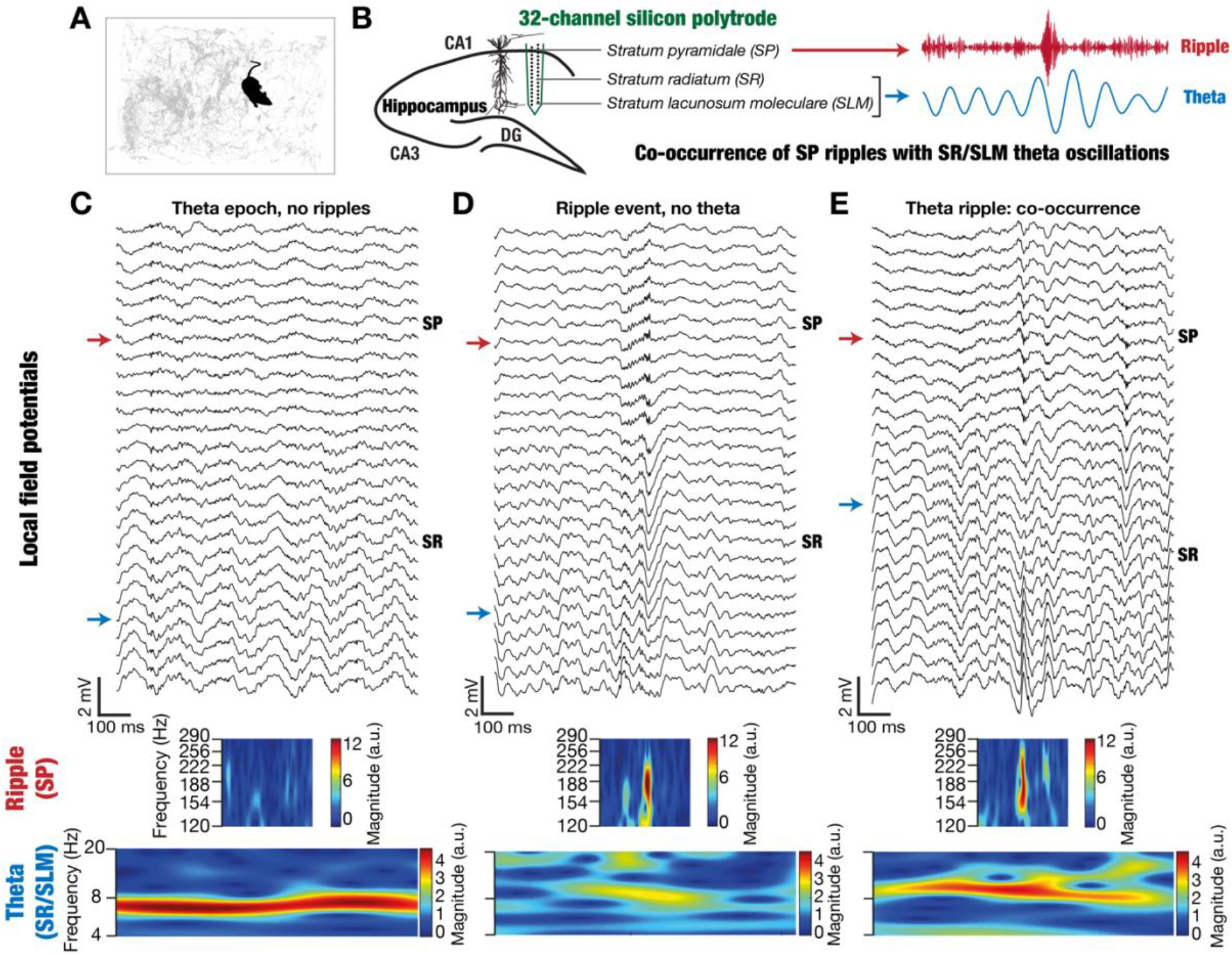
Co-occurrence of theta-frequency oscillations and ripples in a foraging rat. **(A)** Trajectory (gray lines) of a rat in the arena (59 cm × 49 cm) an example recording session. **(B) Left**, Schematic of the hippocampus proper with a single dorsal CA1 pyramidal neuron shown for illustration. Adjacent to the neuron is the sketch of a 32-channel silicon polytrode. The black circles within the electrode are representative of electrode locations relative to the neuron, spanning across the *stratum lacunosum-moleculare* (SLM) to *stratum pyramidale* (SP) stretch. **Right**, ripple-(**top**) and theta-frequency (**bottom**) filtered traces of extracellular voltage recordings simultaneously acquired from the SP and SR/SLM respectively. In these example recordings, theta oscillations in the SR/SLM electrode co-occur with a ripple event in SP electrode. Such ripples in the SP co-occuring with theta-oscillations in the SR/SLM electrode are defined as theta ripples. **(C–E, top)** Laminar profiles of simultaneous LFP recordings during a classical theta epoch specifically showing that the theta amplitude is higher in SR/SLM electrodes compared to SP electrodes **(C)**; a classical sharp wave ripple (SPW-R) complex with sharp waves predominantly visible in the SP-SR electrodes **(D)**; and a co-occurrence event where a ripple in the SP electrodes is embedded within theta-oscillations that are observed prominently in the SR electrodes **(E)**. Red and blue colored arrows refer to the identified SP and SR/SLM *strata* respectively used for time-frequency analysis below. **(C–E, middle)** Wavelet spectrograms of the SP LFP trace shown for the ripple-frequency range (120–250 Hz) indicating the clear prevalence of ripples in panels D and E. **(C–E, bottom)** Wavelet spectrograms of the SR/SLM LFP trace in the theta-frequency range (6–10 Hz) demonstrating strong theta oscillations in panels C and E. Note that the spectrograms were obtained from SP or the SR/SLM LFP trace without any additional filtering in the ripple or the theta frequency ranges. More examples of theta ripples are provided in Fig. S5–S7 and another example of a non-theta ripple is given in Fig. S8.

We identified all valid ripples (total valid ripples: Rat 1: 5293; Rat 2: 8320; Rat 3: 8006) from the SP channel and asked if any of these identified ripples were embedded within epochs of theta oscillations deduced from the SR/SLM LFP. Strikingly, we found several instances of ripple events delineated from the SP signal to be embedded within epochs of theta-frequency oscillations recorded within SR/SLM (Fig. 1*B*). We defined such ripples that occurred within identified theta epochs as *theta* (θ) *ripples* (examples: Fig. 1*E*, Fig. S5–S7), and distinguished them from *non-theta* (*non-*θ) *ripples* that occurred outside of theta epochs (examples: Fig. 1*D*; Fig. S8). The overall proportion of θ ripples across all rats, spanning all behavioral sessions in the open arena, was around 18% (% θ ripples: Rat 1: 444/5293=8.4%; Rat 2: 2430/8320=29.2%; Rat 3: 1325/8006=16.6%). Wavelet scalograms of recordings involving different cross-*strata* combinations — theta oscillations without ripples (Fig. 1*C*), non-theta ripples (Fig. 1*D*), and theta ripples (Fig. 1*E*) — quantitatively confirmed the presence (Fig. 1*C*, Fig. 1*E*) or absence (Fig. 1*D*) of strong theta oscillations in the SR/SLM channel, concurrent with ripple presence (Fig. 1*D–E*) or absence (Fig. 1*C*) in the SP channel.

### A majority of theta ripples occurred during immobile periods of the foraging rat

We divided the behavioral state of the foraging animal into exploratory and immobile periods based on cutoffs on instantaneous velocity and epoch duration. We classified ripples into two categories based on whether they were observed during the exploratory (*exp. ripple*) or the immobile (*imm. ripple*) behavioral phases. Together with the electrophysiological categorization of ripples into theta and non-theta ripples, the behavioral state of the animal provided four distinct categories of ripples: exploratory theta (exp. θ), exploratory non-theta (exp. non-θ), immobile theta (imm. θ), and immobile non-theta (imm. non-θ). We examined the fraction and frequency of occurrence of the four different kinds of ripples based on electrophysiological and behavioral co-occurrences (Fig. 2). Whereas the immobile non-θ ripples constituted a major proportion of all ripples (∼80% across all rats), most of the remaining 20% of ripples were immobile θ ripples. Exploratory ripples during the foraging task were very few, irrespective of whether SP ripples occurred with or without co-occurring SR/SLM theta oscillations (Fig. 2*A*). Notably, across sessions, the frequency of immobile theta ripples was significantly higher than immobile non-θ ripples in rats 2–3, while the two quantities were comparable in Rat 1 (Fig. 2*B*). These observations imply that the probability of observing more immobile θ ripples adjacent to each other is higher than that for the other three kinds of ripples. In two of the rats (Rats 2–3), we repeated our experiments and analyses in another arena, a linear maze (Fig. S9*A*). Our conclusions about the proportions of different ripple subtypes (Fig. S9*B*) and frequencies of their occurrence were consistent across the open arena (Fig. 2) and the linear maze (Fig. S9).

**Figure 2.**
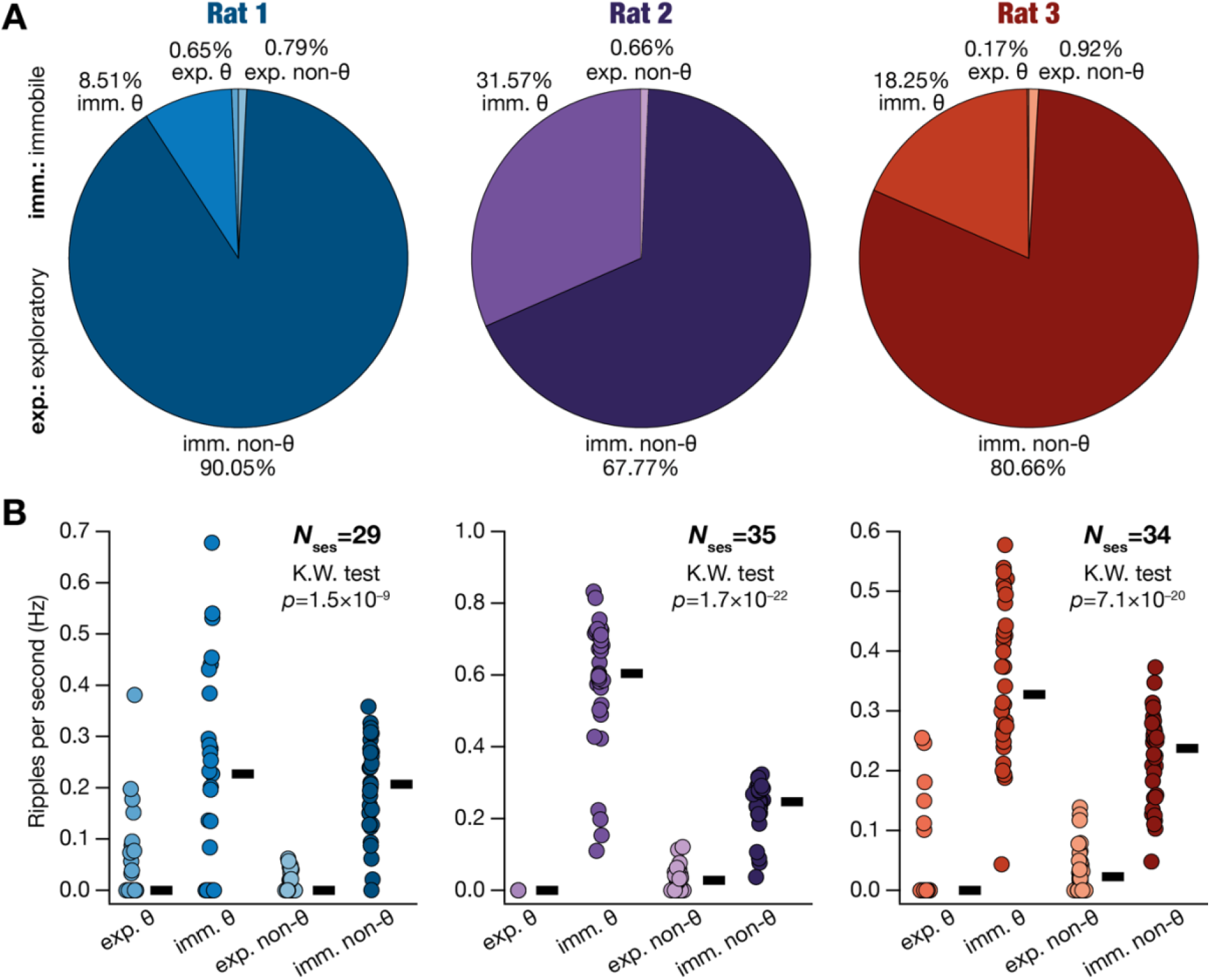
Fraction and frequency of different kinds of ripples. **(A)** Pie charts representing proportions of the four kinds of ripples — exploratory theta (exp. θ); immobile theta (imm. θ); exploratory non-theta (exp. non-θ); and immobile non-theta (imm. non-θ) — for the three rats spanning all recording sessions. **(B)** Frequency of occurrence of ripples for the four ripple kinds, provided for all recorded sessions. The thick black lines represent the respective median values across all sessions. The *p* values computed with the Kruskal Wallis (K.W.) test are provided in the figure for each rat. The group-wise Wilcox rank sum test *p* values: **Rat 1**: exp. θ *vs*. imm. θ: 0.0012, imm. θ *vs*. exp. non-θ: 2.1×10^−4^, exp. non-θ *vs*. imm. non-θ: 7.1×10^−20^, exp. θ *vs*. imm. non-θ: 6.1×10^−7^, exp. θ *vs*. exp. non-θ: 0.3775, imm. θ *vs*. imm. non-θ: 0.85; **Rat 2**: exp. θ *vs*. imm. θ: 1.7×10^−9^, imm. θ *vs*. exp. non-θ: 7.6×10^−13^, exp. non-θ *vs*. imm. non-θ: 3.2×10^−12^, exp. θ *vs*. imm. non-θ: 1.7×10^−9^, exp. θ *vs*. exp. non-θ: 5.9×10^−7^, imm. θ *vs*. imm. non-θ: 1.8×10^−10^; **Rat 3**: exp. θ *vs*. imm. θ: 2.0×10^−10^, imm. θ *vs*. exp. non-θ: 3.1×10^− 12^, exp. non-θ *vs*. imm. non-θ: 8.1×10^−12^, exp. θ *vs*. imm. non-θ: 1.2×10^−8^, exp. θ *vs*. exp. non-θ: 0.032, imm. θ *vs*. imm. non-θ: 5.1×10^−6^.

### Theta and non-theta ripples had comparable characteristics

Are theta ripples distinct from non-theta ripples with reference to the several signature characteristics of ripples? To address this, we computed multiple signature measurements associated with all the detected ripples (maximum frequency, amplitude, duration, power, velocity, SR/SLM theta power during the ripple) and compared them for theta *vs*. non-theta ripples (Fig. 3; Figs. S10–S12). We found all ripple measurements (maximum frequency, amplitude, duration, power) and their distributions to be similar between the theta and the non-theta groups, across all three rats (Fig. 3*A–C*; Figs. S10–S12). As most ripples occurred during immobile periods (velocity < 5 cm/s for at least 1 s), the velocity of the animal during ripple occurrences were also comparable for theta *vs*. non-theta ripples (Figs. S10–S12). These analyses demonstrated that theta-ripples were not intrinsically distinct from the traditional non-theta ripples. As theta power distinguished theta and non-theta ripples, we found the theta power during the occurrence of the ripple to be significantly higher for theta ripples compared to that for non-theta ripples (Figs. S10–S12). Importantly, theta power in the SR/SLM channel during theta ripples (50 ms on either side around the peak of the ripple) was comparable (Fig. 3*D*) with theta power observed during theta epochs (power computed across the entire epoch). Theta and non-theta ripple measurements did not show strong pairwise relationships (Figs. S10–S12), except for the expected strong relationship between theta power and theta amplitude.

**Figure 3.**
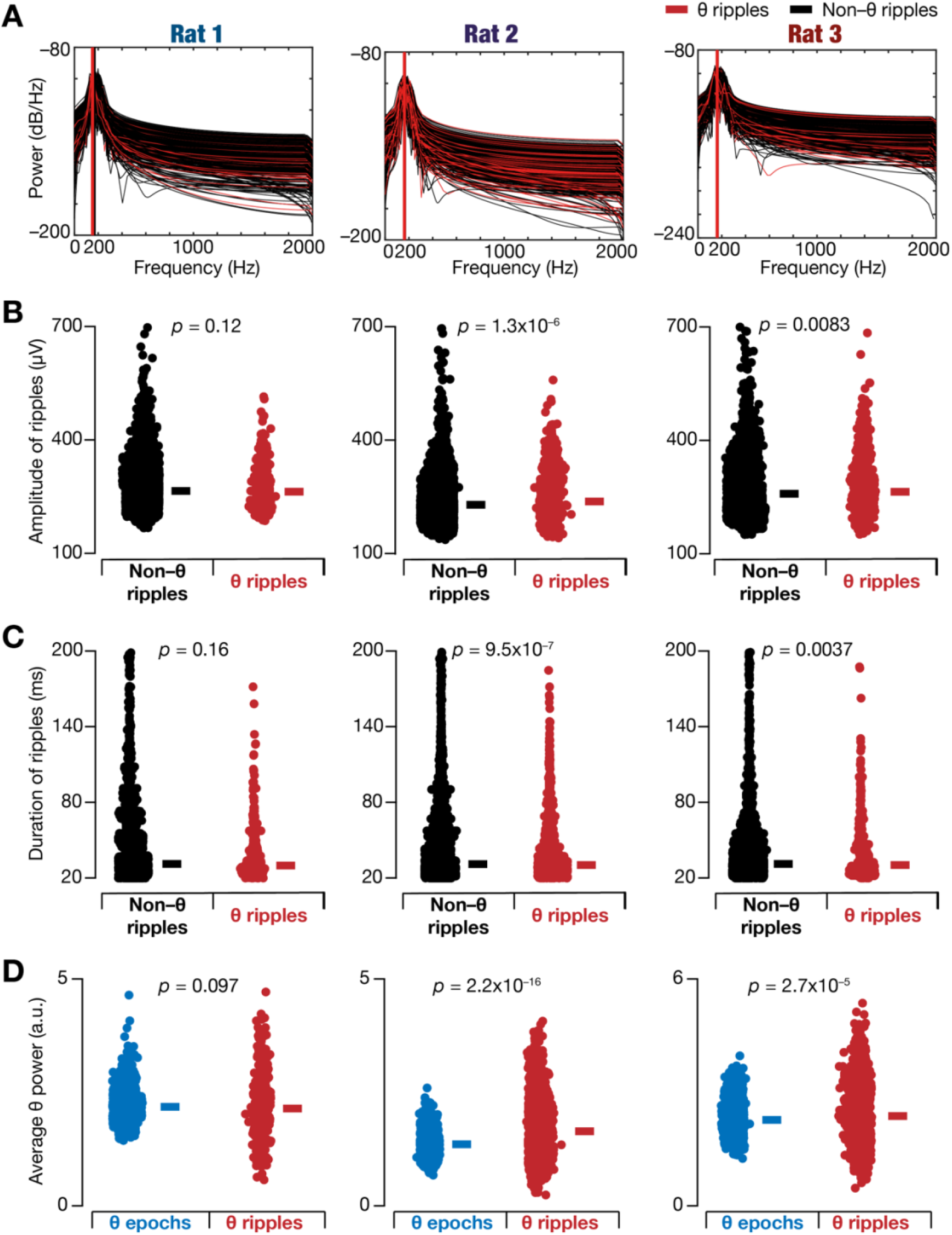
Comparable characteristics of theta and non-theta ripples. **(A–C)** Comparative assessment of the defining characteristics of non-theta and theta ripples, demonstrating the validity of theta ripples. **(A)** Power-spectral density (PSD) of non-theta (black) and theta (red) ripple traces, showing comparable spectral properties of theta and non-theta ripples. Each PSD trace corresponds to an individual ripple. From every session, a random set of 10% of ripples were chosen for this illustration. The vertical lines indicate the average frequency of maximal power for non-theta (black) and theta (red) ripples. **(B–C)** Comparable amplitudes **(B)** and durations **(C)** of non-theta (black) and theta (red) ripples. Each point refers to individual ripples obtained across all sessions in each of the three rats. **(D)** Comparable magnitudes of theta power during classic theta epochs (blue) and during theta-ripple events (50 ms on either side of the peak of the ripple; total 100 ms period in ripple vicinity) shows the validity of theta oscillations observed during theta ripple events. Each point refers to the individual theta epochs or individual ripples across all sessions. *p* values in B–D correspond to Wilcox rank sum test.

Although the overall durations of theta and non-theta ripples were comparable (Fig. 3*C*), we asked if there were differential expression profiles of long *vs*. short ripples (cutoff: 80 ms) within theta epochs (Fig. 4). We found session-to-session and animal-to-animal variability in the proportions of short/long ripples within theta epochs (Fig. 4*A–B*), but the overall proportions of short/long ripples followed the percentage of theta ripples among all identified ripples (Fig. 2*A*). The percentage of theta or non-theta ripples that were long also manifested variability across sessions and animals but were comparable in their ranges (Fig. 4*C–D*). Together, these analyses (Figs. 3–4) demonstrated that theta ripples showed similar characteristics as their non-theta counterparts and that the theta-frequency power during theta ripples and during theta epochs were comparable.

**Figure 4.**
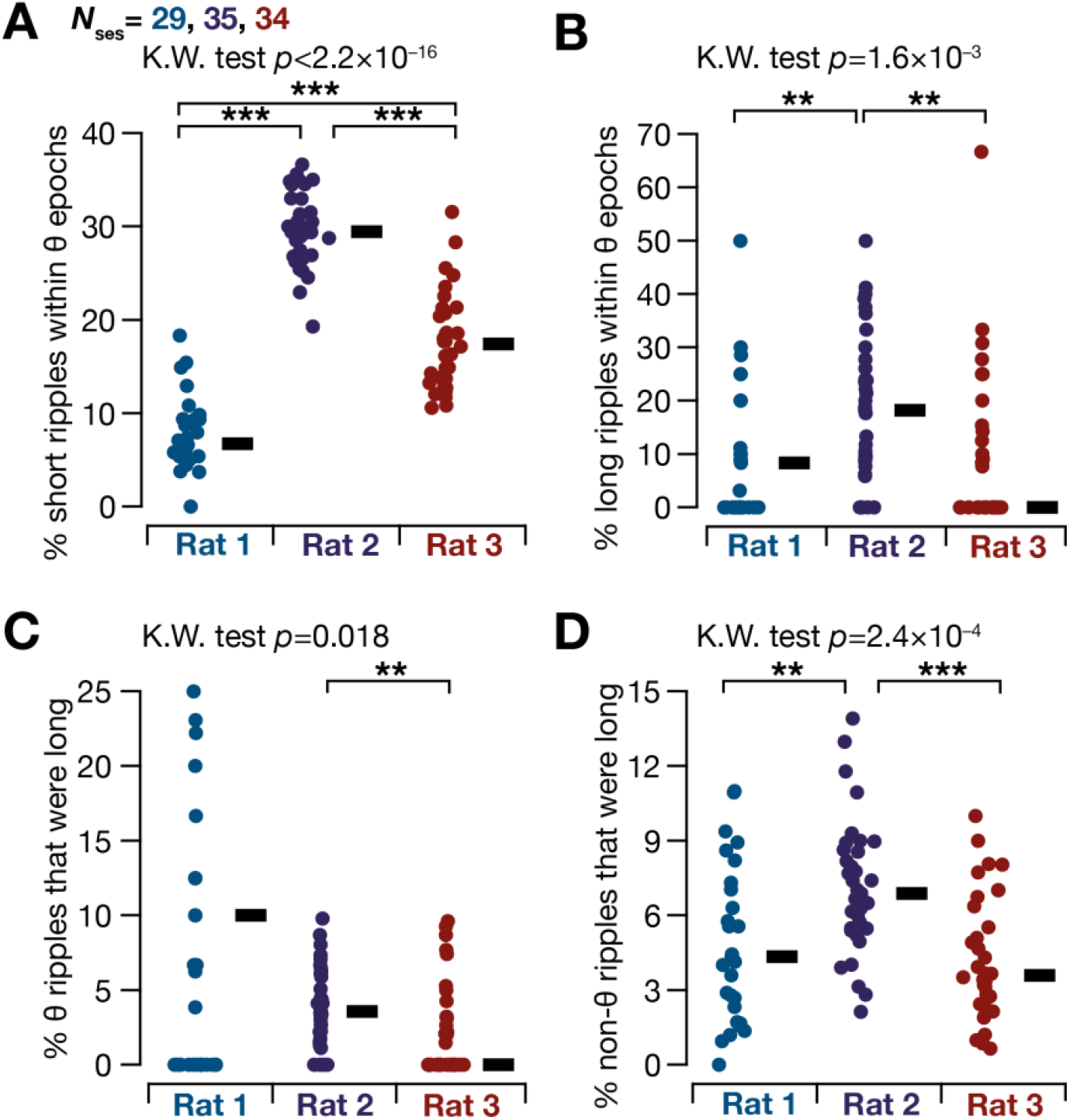
Distributions of ripple duration among theta and non-theta ripples. **(A)** Percentage of short ripples that were detected within theta epochs. **(B)** Percentage of long ripples that were detected within theta epochs. **(C)** Percentage of theta ripples that were of long duration. **(D)** Percentage of non-theta ripples that were of long duration. In all plots the thick black lines indicate the respective median values. In each rat, the percentages were calculated in a session-wise manner and each point corresponds to the respective percentage value computed within a given session. K.W.: Kruskal Wallis. Pairwise significance is provided for Wilcox rank sum test. *: *p<*0.05; **: *p<*0.01; ***: *p<*0.001.

### Theta ripples preferentially occurred within the falling phase of the co-occurring theta oscillations

Do theta ripples detected in the SP occur at specific phases of the theta oscillation concomitantly detected in the SR/SLM? We examined this by computing the phase of the peak of a detected theta ripple in SP relative to the theta oscillations that were simultaneously identified in SR/SLM (example: Fig. S13). Surprisingly, we found that theta ripples had a consistent phase relationship to the concomitant theta oscillations across all rats. Specifically, theta ripples manifested a phase preference predominantly for the fourth quadrant (3π/2–2π phase range) of the associated theta oscillation (Fig. 5). With the trough designated as zero phase, this preference corresponds to the falling part of the theta oscillation in the SR/SLM.

**Figure 5.**
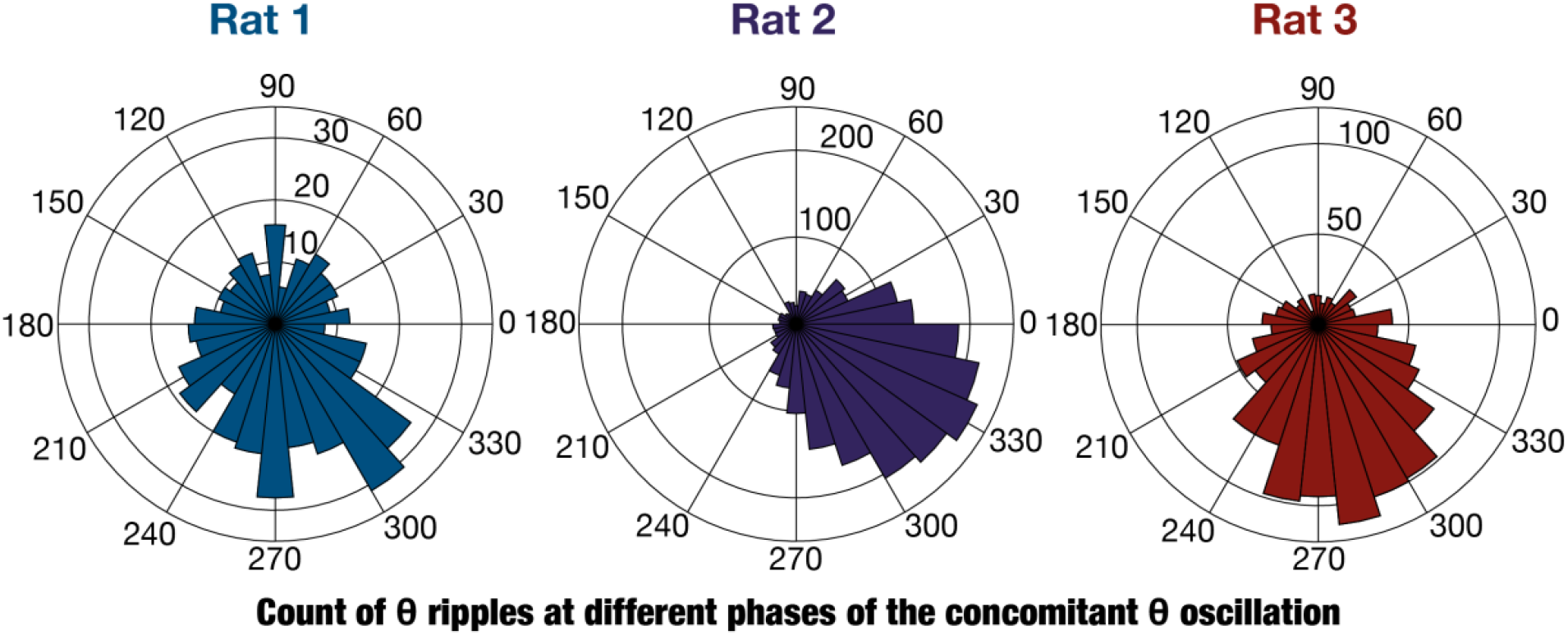
Phase relationship of theta ripples with co-occurring theta oscillations. Polar histogram showing the distribution of phases of the peak of the ripple (SP channel) with respect to the theta oscillations (SR/SLM channel) during theta-ripple events for all rats. It may be noted that phases of theta ripples show a preference for the fourth quadrant of the theta oscillation. Zero degrees represented the trough of the theta oscillation.

### Lower theta power in SP emphasizes the need for simultaneous multi-*strata* recordings in detecting theta ripples

Two important methodological differences in our experimental design and analyses were: (i) the use of simultaneous multi-*strata* recordings; and (ii) analyzing LFPs from two different *strata* for identifying theta oscillations (SR/SLM) and ripples (SP). These experimental and analyses design paradigms were driven by the established differences in expression profiles of theta oscillations and ripples across different *strata* (Buzsaki, 2002; Gordon et al., 2005; Zutshi et al., 2022). Theta oscillations are dominant in the SLM owing to the distal localization of entorhinal cortical inputs and progressively reduced in theta power along the SLM-SR-SP-SO axis (Buzsaki, 2002; Gordon et al., 2005; Zutshi et al., 2022). Ripples, on the other hand, are detected predominantly in the perisomatic *strata*, primarily localized to the SP (Gordon et al., 2005; Buzsáki, 2015; Navas-Olive et al., 2022).

We confirmed the differential power of theta oscillations across CA1 *strata* during exploratory and immobile periods of behavior (Fig. 6, Fig. S14). Specifically, our analyses demonstrated that theta power in individually identified theta epochs was higher in the SLM, with a progressive fall in power along the SLM-SR-SP-SO axis during both immobile and exploratory theta epochs. These analyses confirmed that the theta power during immobile periods (during which most theta ripples were identified; Fig. 2) was comparable (Fig. 6, Fig. S14) to theta power during exploratory periods (to which theta oscillations are traditionally associated with). Thus, the ability to detect theta oscillations and associated theta epochs progressively reduce if SP or SO LFP were employed for the purpose. Consequently, we hypothesized that the number of identified theta ripples should reduce if we employed SP or SO channels as the theta channel, rather than using the SR/SLM LFP that we have been using thus far.

**Figure 6.**
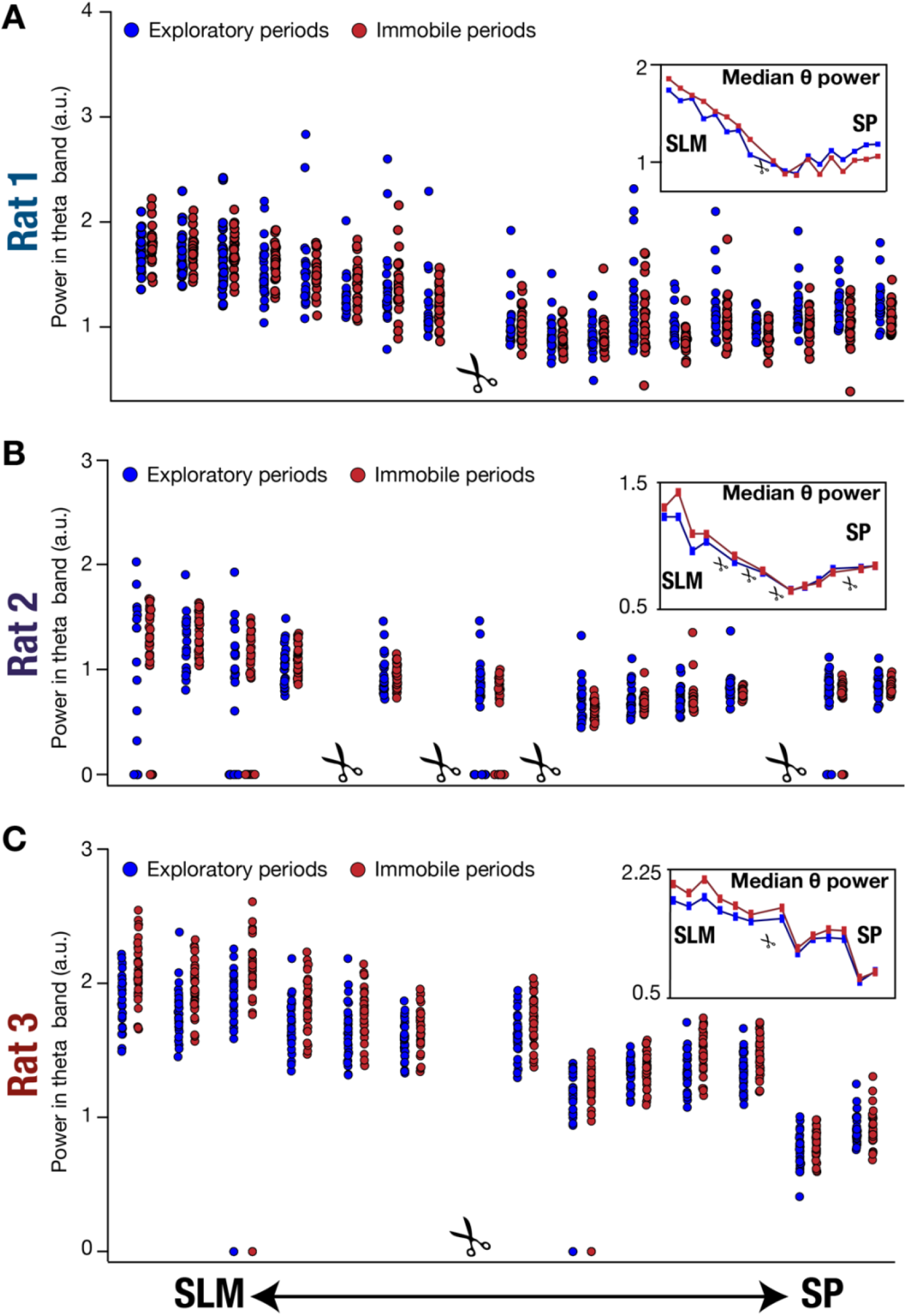
Theta power was highest in SLM with a progressive power reduction through SR to SP during immobile and exploratory periods. Theta power during immobile and exploratory periods obtained from recording channels spanning different *strata* (SLM to SP) for the three rats across all recording sessions. The insets show the *stratum-*dependent median power values computed from the respective points shown in the main plots. A symbol showing a pair of scissors indicates that the specific recording channel was noisy for a majority (90%) of the recorded sessions and was discarded from analyses. Panels A–C represent different rats (rats 1–3). Wilcox rank sum test *p* values for theta power computed from the identified SLM *vs*. SP channel: **Rat 1**: SLM *vs*. SP: 3.18×10^−12^ (exp.), 7.4×10^−12^ (imm.); **Rat 2**: SLM *vs*. SP: 4.91×10^−7^ (exp.), 2.2×10^−16^ (imm.); **Rat 3**: SLM *vs*. SP: 2.2×10^−16^ (exp.), 2.2×10^−16^ (imm.).

To directly test this, we used the ripples detected from the SP channels and asked if the number of theta ripples would change if we used different channels spanning the SLM-SR-SP-SO axis for identifying theta epochs. Specifically, we picked one of the several channels that fell within these axes and used theta-filtered power in that channel to delineate theta epochs. We then asked how many of the ripples identified from the SP channel were within these theta epochs delineated from this chosen channel. We computed this count of theta ripples for each session for each of the three rats, and the whole analysis was performed for all channels along SLM-SR-SP-SO axis (Fig. 7). Strikingly, and consistent with our hypothesis, we found that number of co-occurrences between theta oscillations and ripples were significantly lower if the SP or the SO channels were used for identifying theta epochs (Fig. 7).

**Figure 7.**
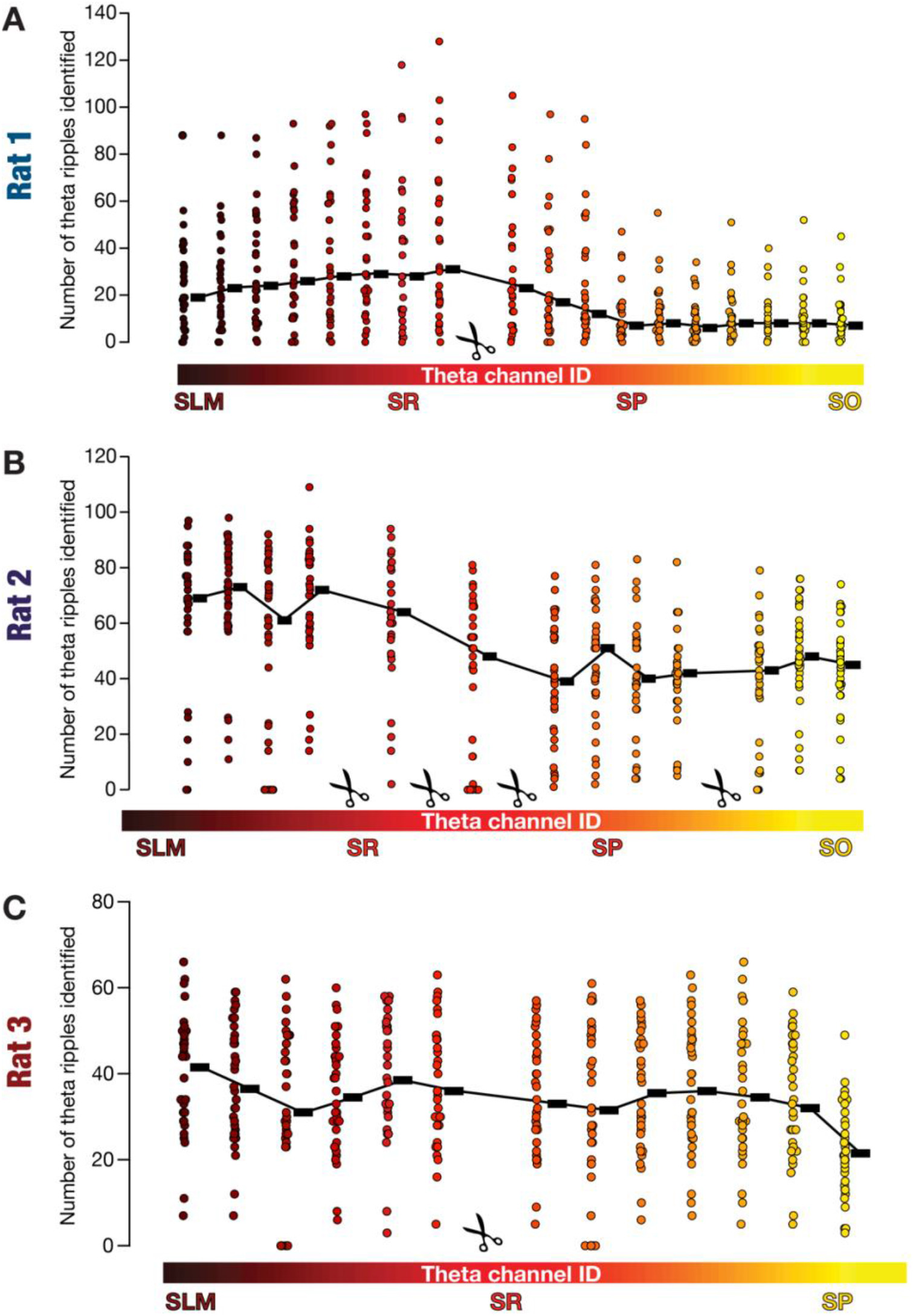
Theta-ripple co-occurrences depended on the channel employed for identifying theta epochs. **(A– C)** The number of theta ripple co-occurrence events are plotted as a function of the identity of the channel from where theta information was derived. Ripples were consistently detected from the identified SP channel for each rat, but theta epoch detection was with reference to each of the different channels spanning all *strata*. The number of theta ripples was computed by counting the number of ripples within theta epochs identified in the chosen theta channel. Each point represents the number of theta ripples identified in each session. The color code indicates the channel that was used to identify theta epochs. The black rectangles indicate the respective median values and are connected by a thin black line. Panels A–C represent different rats (rats 1–3). A symbol showing a pair of scissors indicates that the specific recording channel was noisy for a majority (≥90%) of the recorded sessions and was discarded from analyses. Wilcox rank sum test *p* values for number of theta ripples identified when the theta channel was SLM *vs*. SP: **Rat 1**: SLM *vs*. SP: 2.5×10^−3^; **Rat 2**: SLM *vs*. SP: 1.9×10^−7^; **Rat 3**: SLM *vs*. SP: 1.2 × 10^−5^.

## DISCUSSION

The principal conclusion of this study is the demonstration of the co-occurrence of theta oscillations with ripples by explicitly accounting for the differential expression profile of theta oscillations and ripples across CA1 *strata*. Simultaneous multi-*strata* recordings from foraging rats coupled with an analysis regime that detected these oscillations in the *stratum* where their power is the highest provided the substrate for demonstrating the prevalence of theta ripples. The prevalence of theta ripples opens new avenues of investigation, specifically expanding the potential physiological roles of ripples to encoding regimes.

### Experimental designs and analysis pipelines should explicitly account for cross-*strata* co-occurrence of theta oscillations and ripples

Traditionally, studies recording extracellular signals place extracellular electrodes in the SP with a focus on simultaneously capturing neuronal spiking as well as extracellular field potentials from the cell bodies that are predominantly located in the SP. The focus on the SP also extends to studies that have concomitantly recorded intra- and extra-cellular potentials. However, as later studies have elucidated, it is important to measure extracellular signals from *strata* other than the SP that would reflect dendritic potentials, in addition to deriving individual units by proximal placement of electrodes to the cell bodies (Kamondi et al., 1998b; Gordon et al., 2005; Buzsaki, 2015; Sinha and Narayanan, 2022). The fundamental question posed in this study required us to acquire across-*strata* extracellular potentials to simultaneously investigate oscillations that are known to manifest differential spatial distributions across different *strata*. Such multi-electrode acquisition coupled with cross-*strata* analysis of simultaneously recorded oscillatory patterns in two bands unveiled their coexistence, despite the conventional dichotomy that places them to be exclusively prevalent across different behavioral states.

An important design principle in our study is the unbiased nature of the analyses process where there were no implicit assumptions about the dichotomous expression of the oscillatory patterns (theta and ripples) across different behavioral states or across different *strata*. We analyzed all behavioral states and *strata* to identify theta epochs and placed ripples in different groups based on the behavioral state and co-occurrence with theta oscillations. Our analyses clearly confirmed the differential theta profile that is dominant in the SLM and weak in the SP (Fig. 6). This differential power expression implied that the number of theta ripples that were detected were lesser in number when the SP was used as the theta channel (Fig. 7). These analyses also show that the theta power and the associated *strata*-dependence during immobile periods in the SLM channel was comparable to SLM theta power during awake behaving periods (Fig. 6), also arguing against different sources of theta observed during immobile *vs*. exploratory periods.

Together, our study presents a fundamental design requirement for any analyses pipeline as the need to avoid any assumption regarding one-to-one mapping between behavioral states and oscillatory patterns. Explicitly, the analyses of theta oscillations and ripples must involve an unbiased analysis pipeline of simultaneously recorded data from multiple *strata*. The absence of multi-*strata* data or the presence of biased assumptions about the exclusive expression of oscillatory bands (*e*.*g*., theta band oscillations are observed only in the exploratory periods where the velocity is higher than a large threshold) would result in biased conclusions about the co-occurrence of theta oscillations and ripples. Future studies should not assume the prevalence of theta oscillations to mean exploratory behavior or vice versa but should instead explicitly account for the differential power profiles of theta-oscillations (predominantly SLM) and ripples (predominantly SP) while analyzing behavioral correlates of these oscillatory patterns.

### Implications for the co-occurrence of theta oscillations and ripples: Future directions

The reported cross-*strata* co-occurrence of theta oscillations with ripples in the CA1 subregion has important behavioral and physiological implications apart from providing several critical avenues for future explorations. The primary implication for the co-occurrence of theta oscillations and ripples is the need to reassess the dichotomy, driven primarily by historic single-*strata* recordings, of theta oscillations and ripples being exclusively associated with preparatory and consummatory behaviors. Such reassessment would also stipulate expansion of the purported roles of ripple-frequency oscillations to theta epochs as well, especially with reference to replay and preplay of neuronal firing sequences within ripples during sleep and awake behavioral states (Foster and Wilson, 2006; O’Neill et al., 2006; Csicsvari et al., 2007; Diba and Buzsaki, 2007; Jadhav et al., 2012; Nokia et al., 2012; Singer et al., 2013; Fernandez-Ruiz et al., 2019). Specifically, sharp wave ripples have been traditionally implicated in consolidation as well as retrieval of memory (Buzsaki, 1989; Qin et al., 1997; Buzsaki, 1998; Girardeau et al., 2009; Carr et al., 2011; Girardeau and Zugaro, 2011; Buzsáki, 2015; van de Ven et al., 2016; Roux et al., 2017; Wu et al., 2017; Joo and Frank, 2018). Our analyses showing co-expression of theta oscillations with ripples add to the growing interest in a role for ripple-frequency oscillations in other awake behavior including navigational planning, learning, and memory-guided decision making (Foster and Wilson, 2006; O’Neill et al., 2006; Csicsvari et al., 2007; Diba and Buzsaki, 2007; Jadhav et al., 2012; Nokia et al., 2012; Singer et al., 2013; Fernandez-Ruiz et al., 2019).

Awake sharp wave ripples and associated replays have been implicated in several functions (Gupta et al., 2010; Jadhav et al., 2012; Buzsaki, 2015; Papale et al., 2016; Tang et al., 2017; Fernandez-Ruiz et al., 2019) with lines of evidence for a relationship of ripple-related activity to task demands (Olafsdottir et al., 2017). In addition, there is evidence for co-occurrence of theta oscillations and ripples during exploratory periods (O’Neill et al., 2006; Csicsvari et al., 2007) and REM sleep (see (Buzsaki, 2015), Fig. 5 for an example of SPW-R complexes embedded in a stream of theta waves). Importantly, exploratory sharp wave ripples are prevalent in the primate nervous system during visual exploration (Leonard et al., 2015; Leonard and Hoffman, 2017) and there are differences between awake and sleep sharp wave ripples (O’Neill et al., 2006; Csicsvari et al., 2007; Csicsvari and Dupret, 2014). Therefore, an important next step to our report of cross-*strata* co-occurrence of theta-frequency oscillations with ripples during a foraging task is the need to assess such co-occurrence across different behavioral tasks and sleep states. Future studies should explore the prevalence and characteristics of theta ripples across different behavioral states, explicitly assessing cross-dependencies on the task, the novelty of the environment, the state of the animal, pathological conditions, neuromodulatory influences, and within-task switches. Such analyses should employ simultaneous multi-*strata* recordings coupled with unbiased analyses pipelines that evaluate cross-*strata* signatures of ripples and theta oscillations. In this context, addressing the temporal dynamics of spikes associated with theta ripples across different neurons and associated replays and preplays, especially with specific reference to concurrent theta oscillations in different *strata*, across distinct behavioral states constitutes an important requirement. These questions should be addressed both from the intra-as well as extra-cellular perspectives with somato-dendritic voltage dynamics and cross-*strata* LFPs assessed simultaneously.

The analyses above could expand the potential roles of ripple-frequency oscillations to span the continuum of encoding as well as consolidation, through potential interactions with theta oscillations in the process. Specifically, it is possible that encoding and consolidation processes need not be entirely separated in time and that very brief instances of awake immobility where theta oscillations are also prevalent could potentially engage themselves in both phenomena. In this context, the plateau potentials which have been strongly linked to place-cell formation as well as to sharp waves provide strong links to the processes of consolidation as well as encoding (Dragoi et al., 1999; O’Neill et al., 2006; O’Neill et al., 2008; Vandecasteele et al., 2014; Bittner et al., 2015; Buzsaki, 2015; Bittner et al., 2017; Roux et al., 2017; Fernandez-Ruiz et al., 2019; Magee and Grienberger, 2020; Jarzebowski et al., 2021; McKenzie et al., 2021; Valero et al., 2022). The prevalence of plateau potentials during theta oscillations and their roles in place cell formation (Bittner et al., 2015; Bittner et al., 2017) constitutes an important link, and should be explored with coupled intra- and extra-cellular recordings along the somato-dendritic axis. The role of plateau potentials associated with theta ripples in plasticity as well as place cell emergence, and the question of how they affect hippocampal-cortical interactions during theta *vs*. non-theta ripples are important lines of investigation. Within the realm of plasticity, given the cross-*strata* expression of ripples and theta oscillations, future studies could also explore plasticity associated with cross-pathway interactions between entorhinal and CA3 inputs and their potential role in novelty detection and place cell formation (Takahashi and Magee, 2009; Basu et al., 2016; Bittner et al., 2017; Magee and Grienberger, 2020).

In addition to exploring the prevalence and the physiological roles of co-occurring theta ripples across behavioral states, interventional approaches could be employed to gather direct lines of evidence for the specific roles of theta ripples. For instance, simultaneous detection of SR/SLM theta and ripples in SP, coupled with manipulation of the properties or the prevalence (Dragoi et al., 1999; Vandecasteele et al., 2014; Talakoub et al., 2016; van de Ven et al., 2016; Roux et al., 2017; Valero et al., 2017; Fernandez-Ruiz et al., 2019; Karimi Abadchi et al., 2020; Aleman-Zapata et al., 2021; Jarzebowski et al., 2021; Tingley et al., 2021) of theta ripples (and/or associated theta oscillations) could provide insights about theta ripples in memory encoding, consolidation, and storage. The link between the medial septum, a source for theta oscillations to the hippocampal formation (Lee et al., 1994; Buzsaki, 2002), and a metabolic functional role for sharp-wave ripples (Tingley et al., 2021) provides an additional avenue for intervention in exploring the physiological implications of theta ripples. Such interventional experiments should be performed with simultaneous multi-*strata* recordings towards reassessing interactions between medial septum and hippocampal SPW-Rs (Dragoi et al., 1999; Vandecasteele et al., 2014; Jarzebowski et al., 2021), explicitly accounting for theta ripples.

A natural extension for the analyses presented here would be the question of whether such cross-*strata* co-occurrence manifests in the CA3 and the CA2, which play critical roles in the emergence of ripples (Ylinen et al., 1995; Chrobak and Buzsaki, 1996; Buzsaki, 2015) apart from manifesting theta oscillations (Buzsaki, 2002; Goutagny et al., 2009; Colgin, 2013). Furthermore, although our recordings and analyses were limited to the dorsal hippocampus, it is critical to assess co-occurrences with simultaneous multi-*strata* recordings along the dorsoventral and proximo-distal axes of the hippocampus involving brain-wide interactions during such events (Nitzan et al., 2022). Importantly, future studies analyses should assess the intracellular counterparts of theta *vs*. non-theta ripples in the somatic and the dendritic compartments of identified deep *vs*. superficial neurons. It is critical to assess this in a location-dependent fashion as dendritic plateaus might not manifest as somatic depolarization owing to the differential innervation profiles of perisomatic inhibitory neurons onto deep *vs*. superficial neurons (Lee et al., 2014; Valero et al., 2015; Sharif et al., 2021). Thus, simultaneous dendritic recordings could probe the presence of intracellular dendritic plateau potentials and intracellular dendritic theta oscillations during theta ripples identified using multi-*strata* recordings. While these are technically challenging experiments, the availability of multi-electrode recording probes along with high-resolution voltage imaging and other recording techniques in awake behaving animals make such analyses feasible (Giocomo, 2015; Abdelfattah et al., 2019; Steinmetz et al., 2021; Valero et al., 2022; Zong et al., 2022). Such analyses could also assess the role of neurons with basal dendrites that act as axons during theta *vs*. non-theta ripples (Hodapp et al., 2022).

## Acknowledgements

The authors thank members of the cellular neurophysiology laboratory for helpful discussions and for comments on a draft of this manuscript. The authors thank Dr. Sachin Deshmukh, Indraja Jakhalekar, Sriram Narayanan, and Dr. Sunandha Srikanth for helpful advice and feedback through different stages of the study. The authors thank Dr. Liset de la Prida for analysis advice and helpful comments. This work was supported by the DBT-Wellcome Trust India Alliance (Senior fellowship to R. N.: IA/S/16/2/502727), the University Grants Commission (R. B.), and the ministry of education (P. S.).

## Notes

### Competing Interest Statement

The authors have declared no competing interest.

