## Supplementary Figures S1-S14 and Supplementary Table S1 for "Cross-*strata* co-occurrence of ripples with theta-frequency oscillations in the hippocampus of foraging rats"

### Supplementary Material

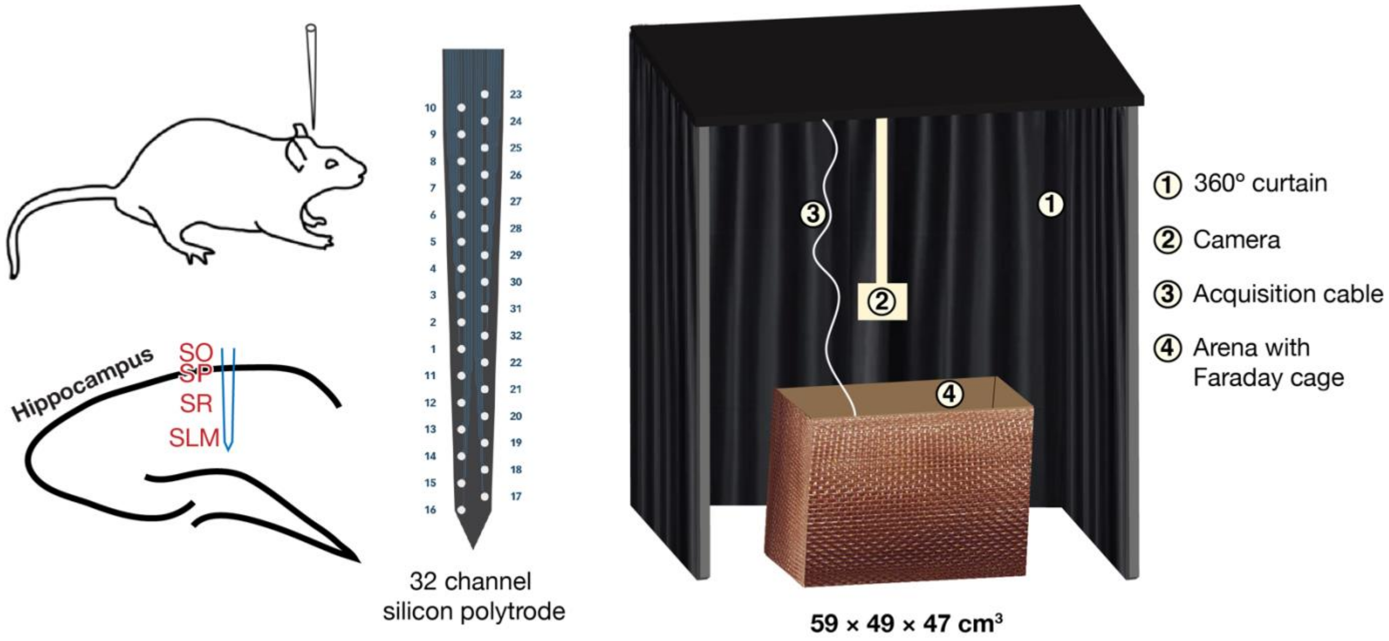

**Supplementary Figure S1. Electrode details and behavioral setup.** *Left*, Stereotaxis apparatus was used to implant electrodes into the dorsal hippocampus such that the silicon polytrode spans the different *strata* of the CA1 region. SO: *Stratum Oriens*. SP: *Stratum Pyramidale*. SR: *Stratum Radiatum*. SLM: *Stratum Lacunosum-Moleculare*. *Middle*, Pictorial representation of the 32-channel silicon polytrode employed for recordings. The numbers correspond to the order of the channels. *Right*, Behavioral setup. The arena was a cardboard box. Behavioral recordings were performed by a ceiling-mounted camera. The entire setup was enclosed within black curtains spanning all sides.

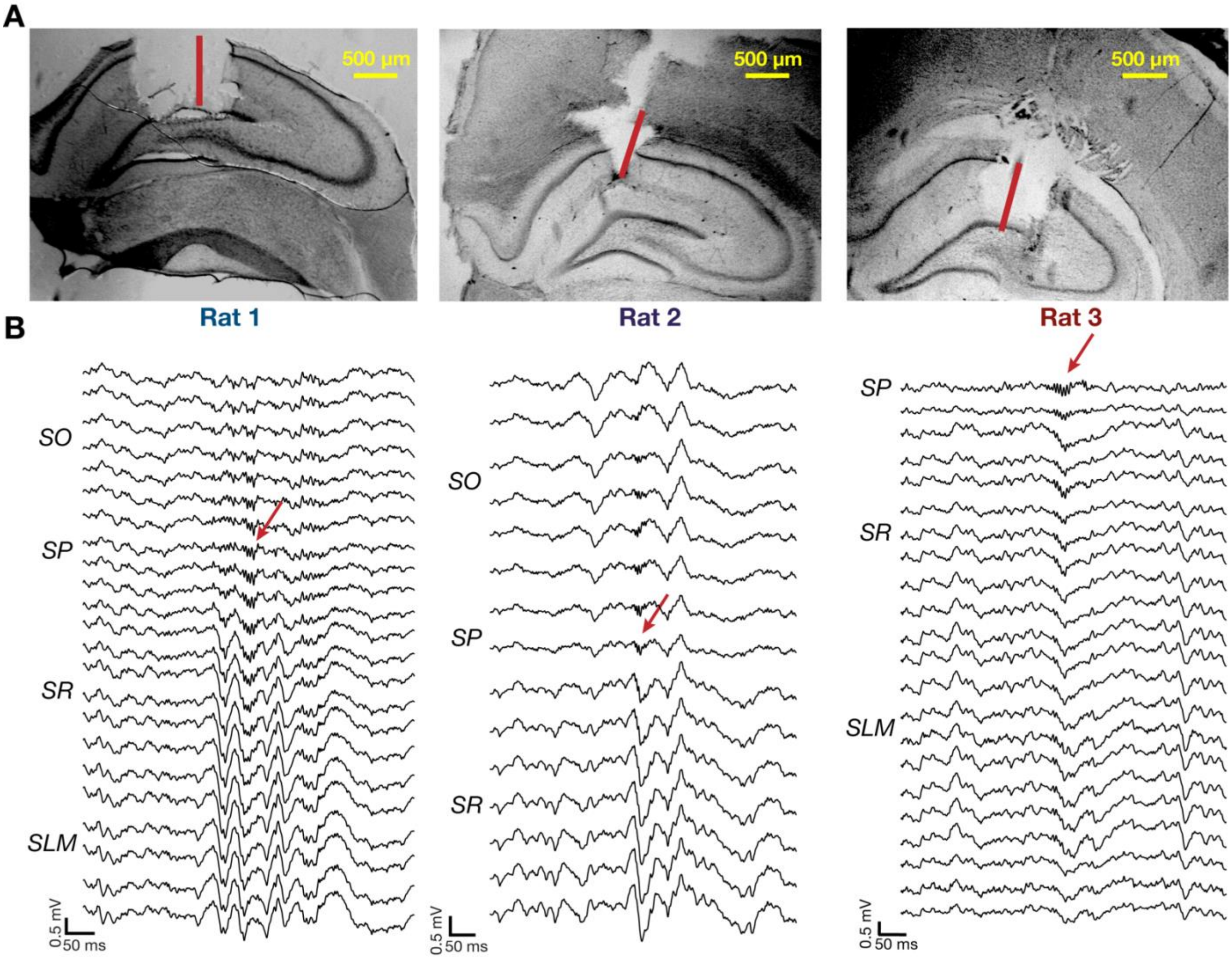

**Supplementary Figure S2. Histology and *strata* identification within CA1. (A)** Grayscale images of the lesioned location and track in the hippocampus of three different rats (left to right). The red bars indicate the inferred location of the 800 μm electrode region of the implanted silicon polytrode based on electrophysiological signatures. **(B)** Laminar profiles showing the local field potentials (0.5 – 300 Hz) recorded in all non-noisy channels from the three rats (left to right). The identity of the *strata* is specified on the left of the laminar profile. Ripples were localized within the SP (marked with a red arrow) and sharp waves are detected within the SR–SLM region. SO: *Stratum Oriens*. SP: *Stratum Pyramidale*. SR: *Stratum Radiatum*. SLM: *Stratum Lacunosum-Moleculare*.

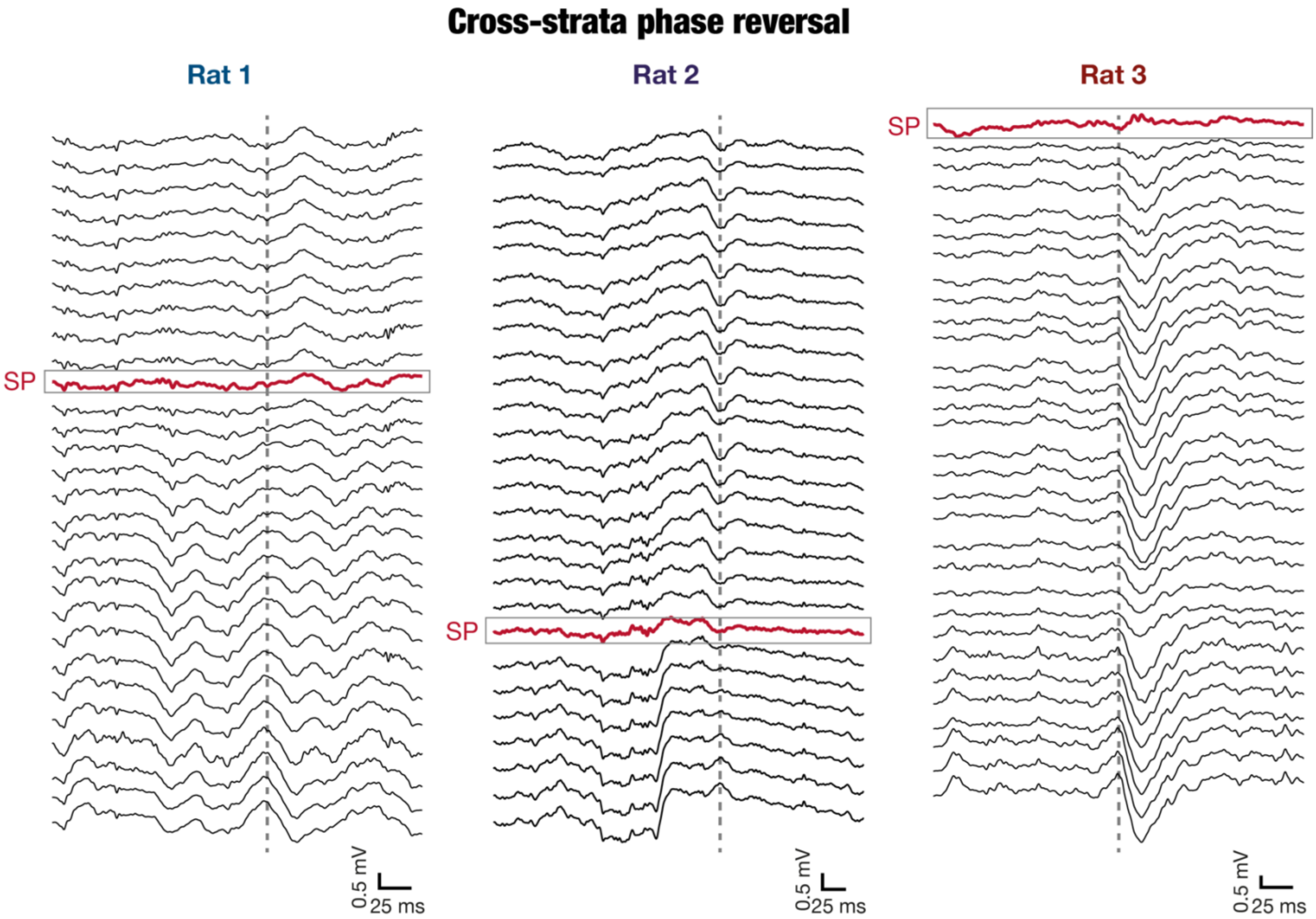

**Supplementary Figure S3. Phase reversal indicating the identity of the *stratum pyramidale* channel.** Laminar profiles showing local field potentials of the three rats (left to right) to highlight the phase reversal occurring at the SP. The grey dashed lines indicate phase reversal, wherein troughs in the SR and SLM layers grow progressively flatter and turn into a peak (or vice versa) as we go up towards the SO. Such a transition at the marked (red colored trace) location further corroborates the identity of the channel that corresponds to the SP. These marked SP channels can be compared with the laminar profiles in Fig. S1 showing the same using sharp wave ripple (SPW-R) complexes.

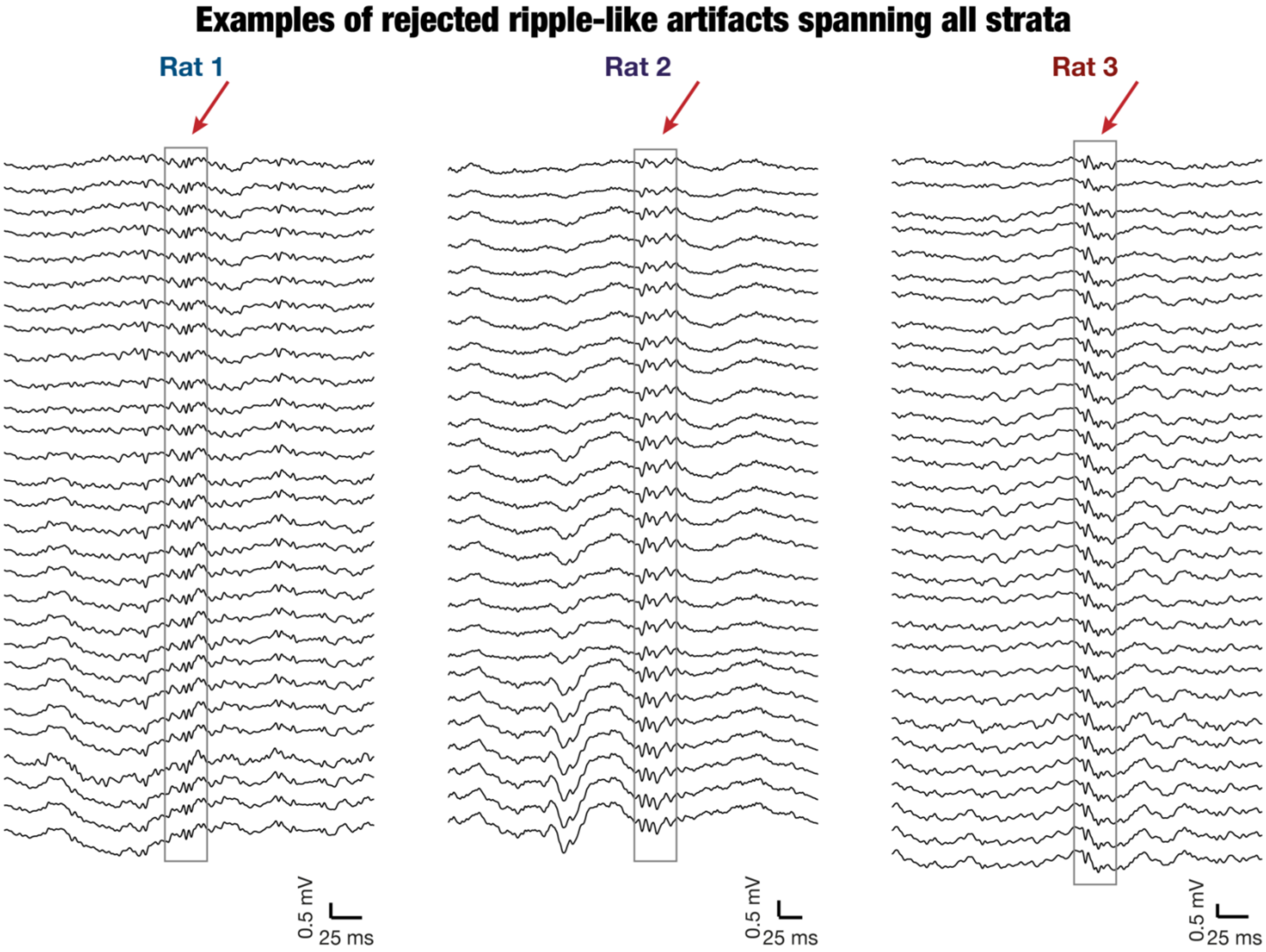

**Supplementary Figure S4. Ripple validation procedure.** Laminar profiles of local field potentials recorded from the three rats (left to right), indicating an event that was initially identified to be a ripple event and then discarded, based on the validation methodology used (see methods). Ripple events, characterized by a specific range of frequency, duration, and inter-ripple interval, were first identified within all recorded channels independently, irrespective of the *strata* they occupied. Events that were detected in more than 10 channels and within a timescale of 10 ms on either side were discarded. Shown above are examples of events that were discarded through this procedure in each of the three rats.

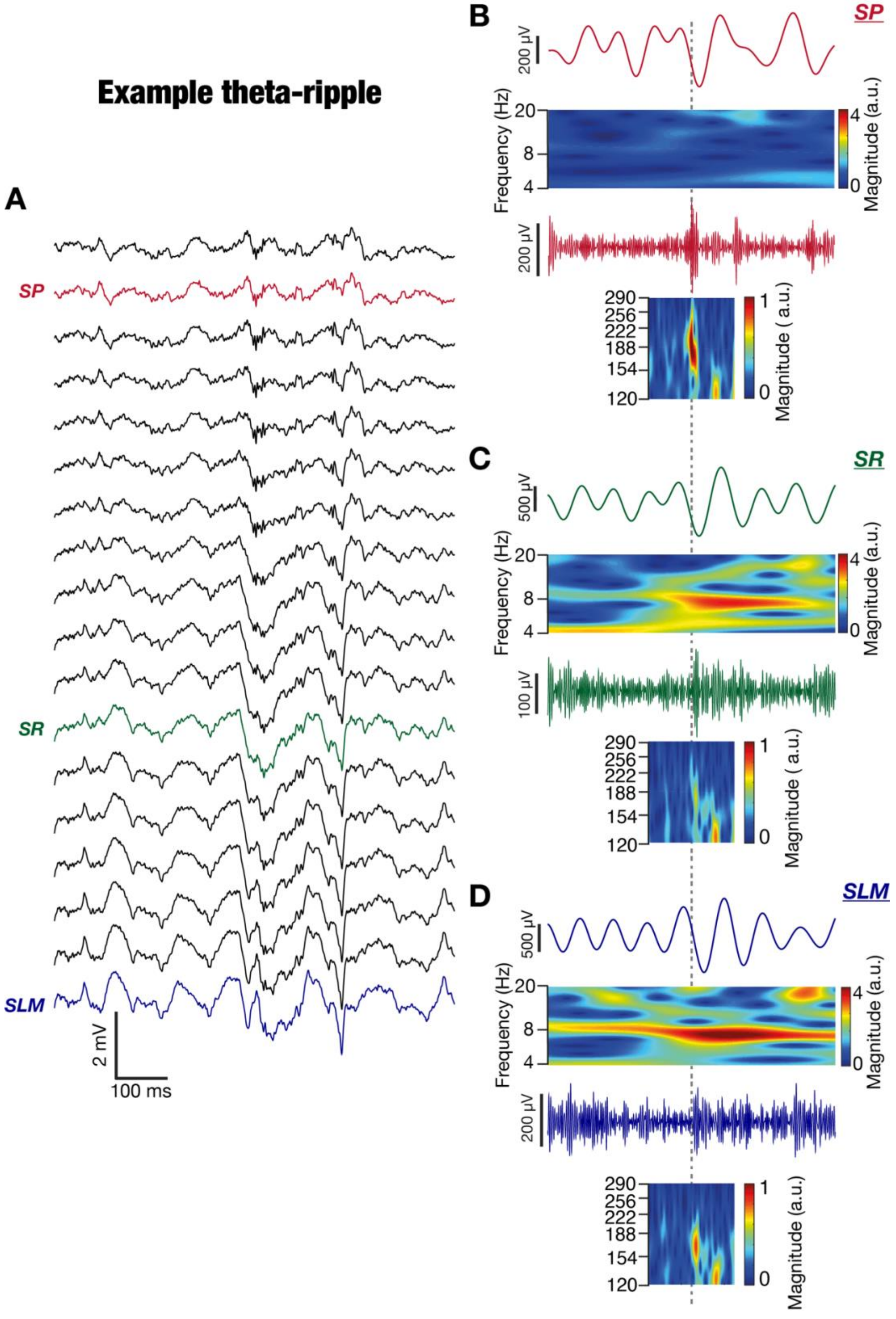

**Supplementary Figure S5. Theta ripple co-occurrence example 1 from rat 1: *Strata*-wise theta- and ripple-frequency spectrograms. (A)** Laminar profile of local field potentials recorded during a theta ripple co-occurrence event with red, green and blue traces indicating SP, SR and SLM respectively. **(B–D)** Theta- (top) and ripple- (bottom) frequency filtered traces and wavelet scalograms for the three identified *strata*. It may be noted that the theta oscillation was strongest in the SLM, whereas the ripple was prominently observed in the SP. The color scale for both theta- and ripple-frequency wavelet scalograms are matched across panels. All scalograms were on unfiltered LFP traces.

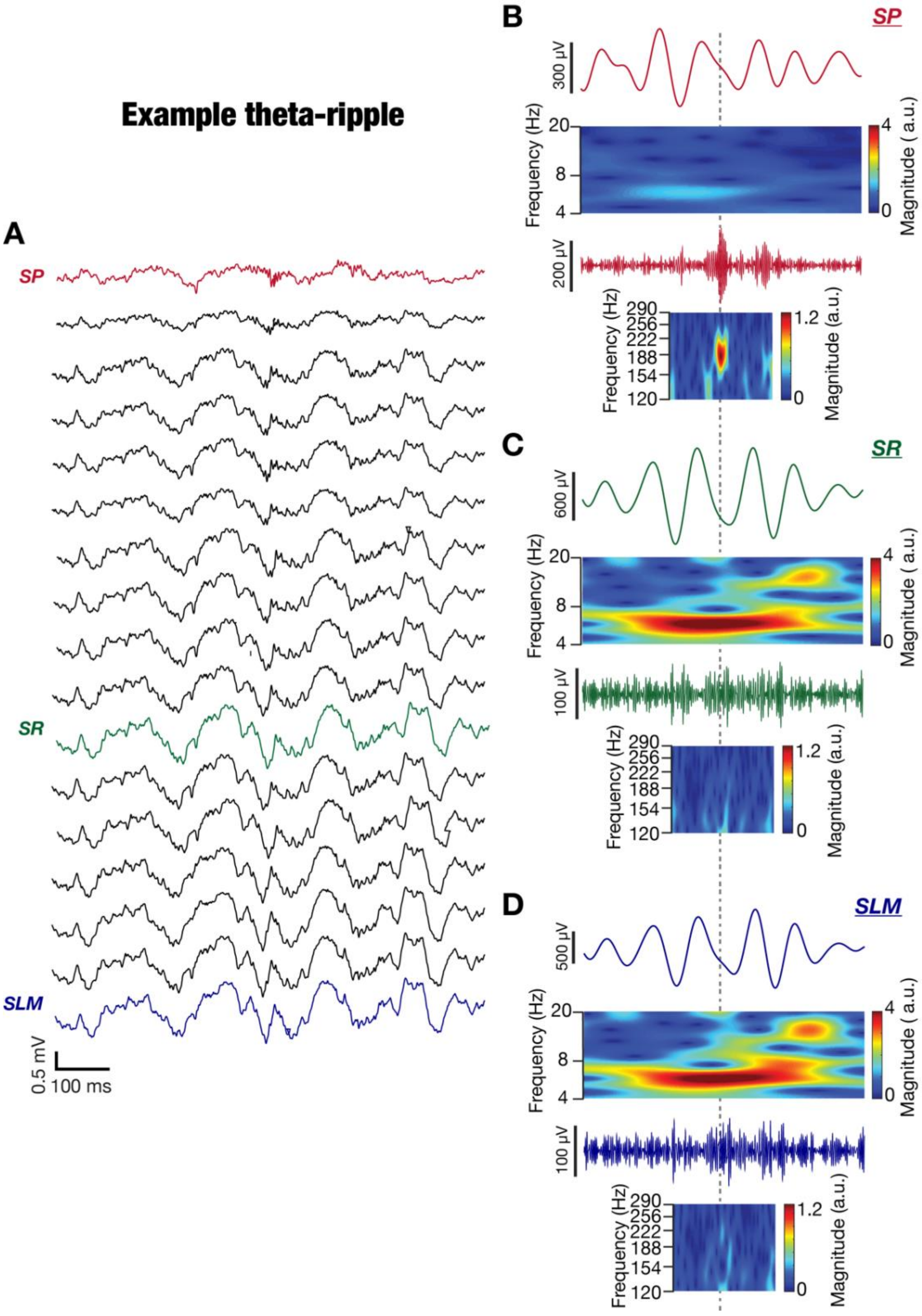

**Supplementary Figure S6. Theta ripple co-occurrence example 2 from rat 3: *Strata*-wise theta- and ripple-frequency spectrograms. (A)** Laminar profile of local field potentials recorded during a theta ripple co-occurrence event with red, green and blue traces indicating SP, SR and SLM respectively. **(B–D)** Theta- (top) and ripple- (bottom) frequency filtered traces and wavelet scalograms for the three identified *strata*. It may be noted that the theta oscillation was strongest in the SLM, whereas the ripple was prominently observed in the SP. The color scale for both theta- and ripple-frequency wavelet scalograms are matched across panels. All scalograms were on unfiltered LFP traces.

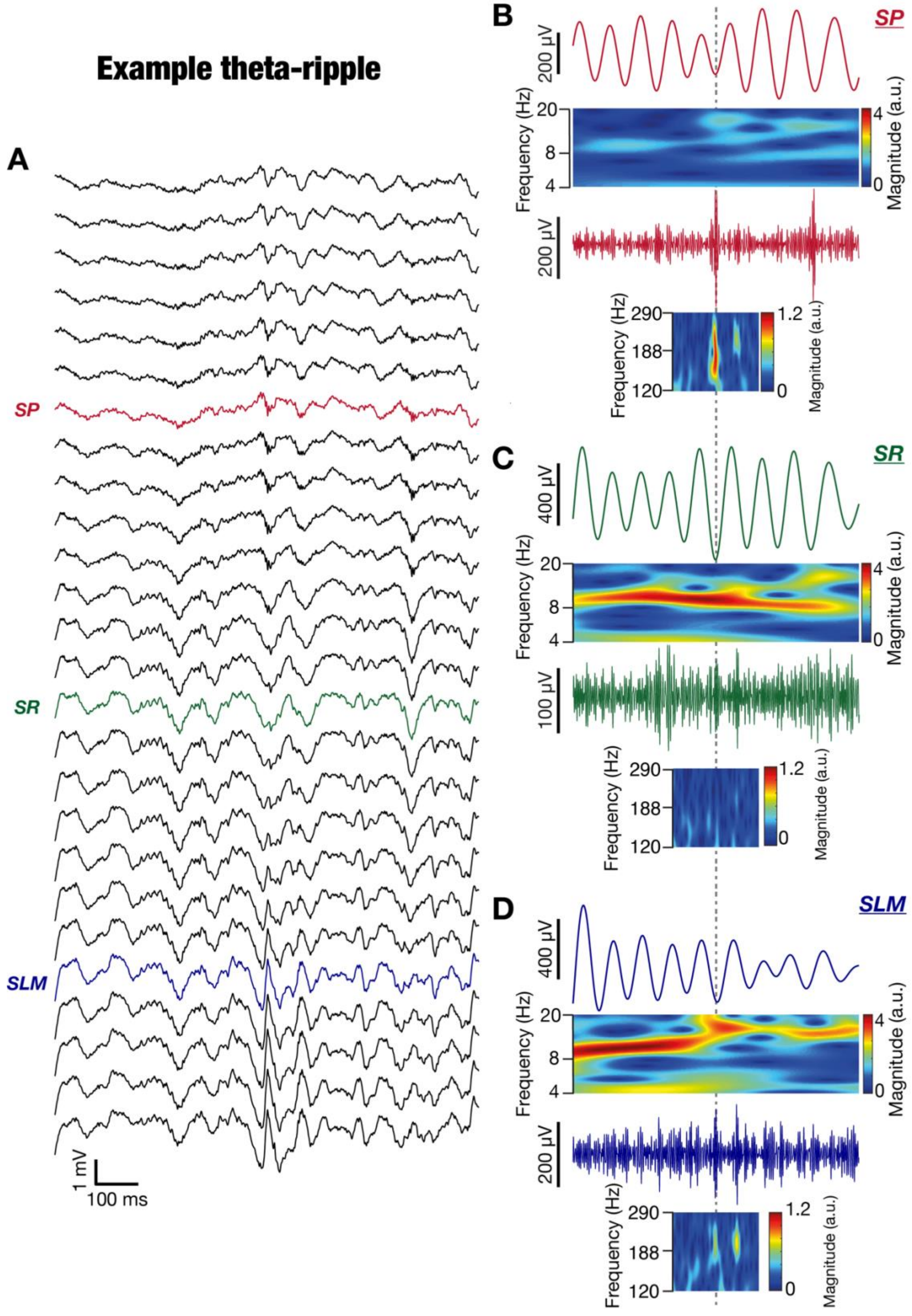

**Supplementary Figure S7. Theta ripple co-occurrence example 3 from rat 1: *Strata*-wise theta- and ripple-frequency spectrograms. (A)** Laminar profile of local field potentials recorded during a theta ripple co-occurrence event with red, green, and blue traces indicating SP, SR and SLM respectively. **(B–D)** Theta- (top) and ripple- (bottom) frequency filtered traces and wavelet scalograms for the three identified *strata*. It may be noted that the theta oscillation was strongest in the SLM, whereas the ripple was prominently observed in the SP. The color scale for both theta- and ripple-frequency wavelet scalograms are matched across panels. All scalograms were on unfiltered LFP traces.

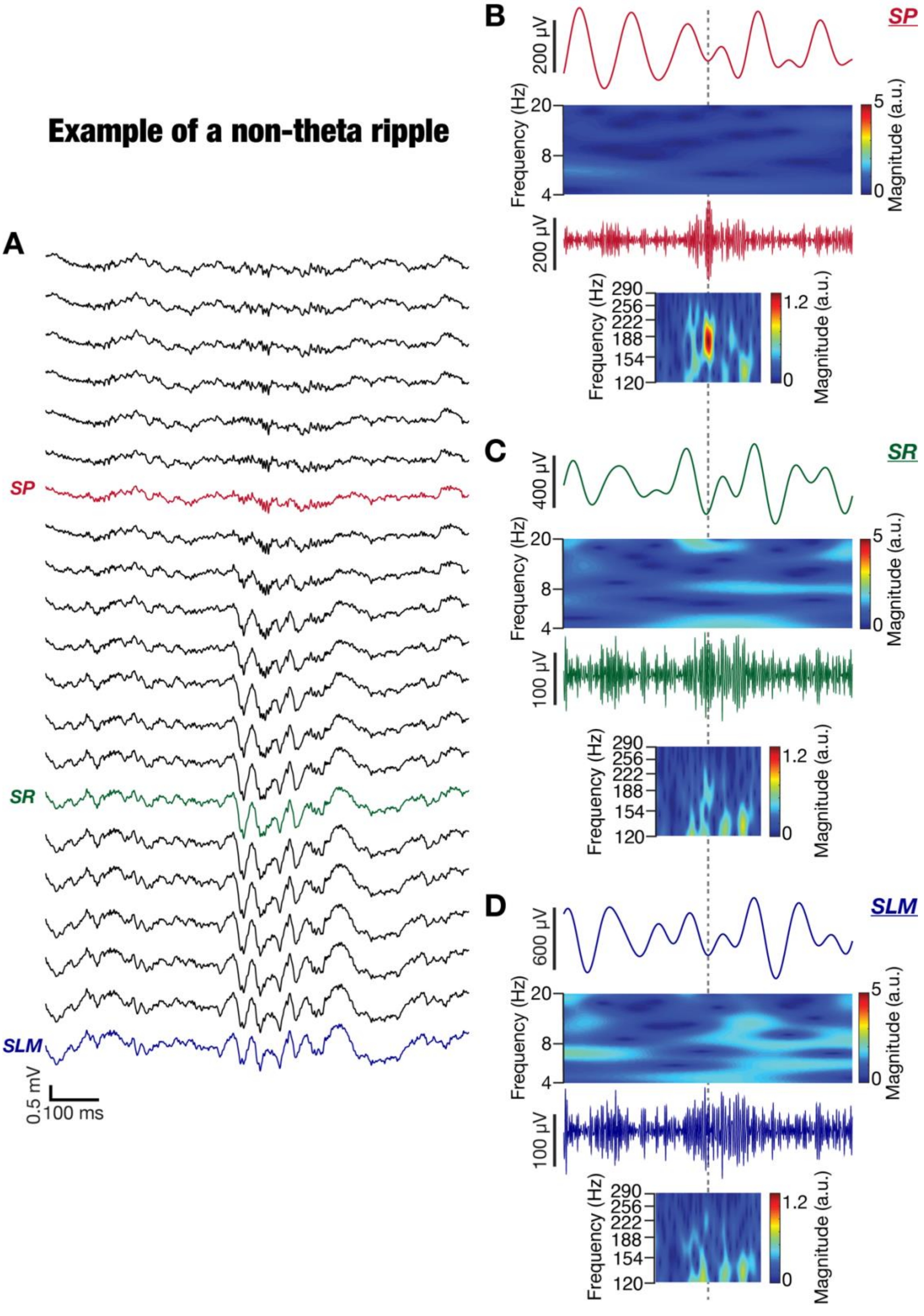

**Supplementary Figure S8. Non-Theta ripple example from rat 1: *Strata*-wise theta- and ripple-frequency spectrograms. (A)** Laminar profile of local field potentials recorded during a sharp wave ripple (SPW-R) event with red, green, and blue traces indicating SP, SR and SLM respectively. **(B–D)** Theta- (top) and ripple- (bottom) frequency filtered traces and wavelet scalograms for the three identified *strata*. It may be noted that the theta oscillations were not observed in any of the channels. The ripple was prominently observed in the SP. The color scale for both theta- and ripple-frequency wavelet scalograms are matched across panels. All scalograms were on unfiltered LFP traces.

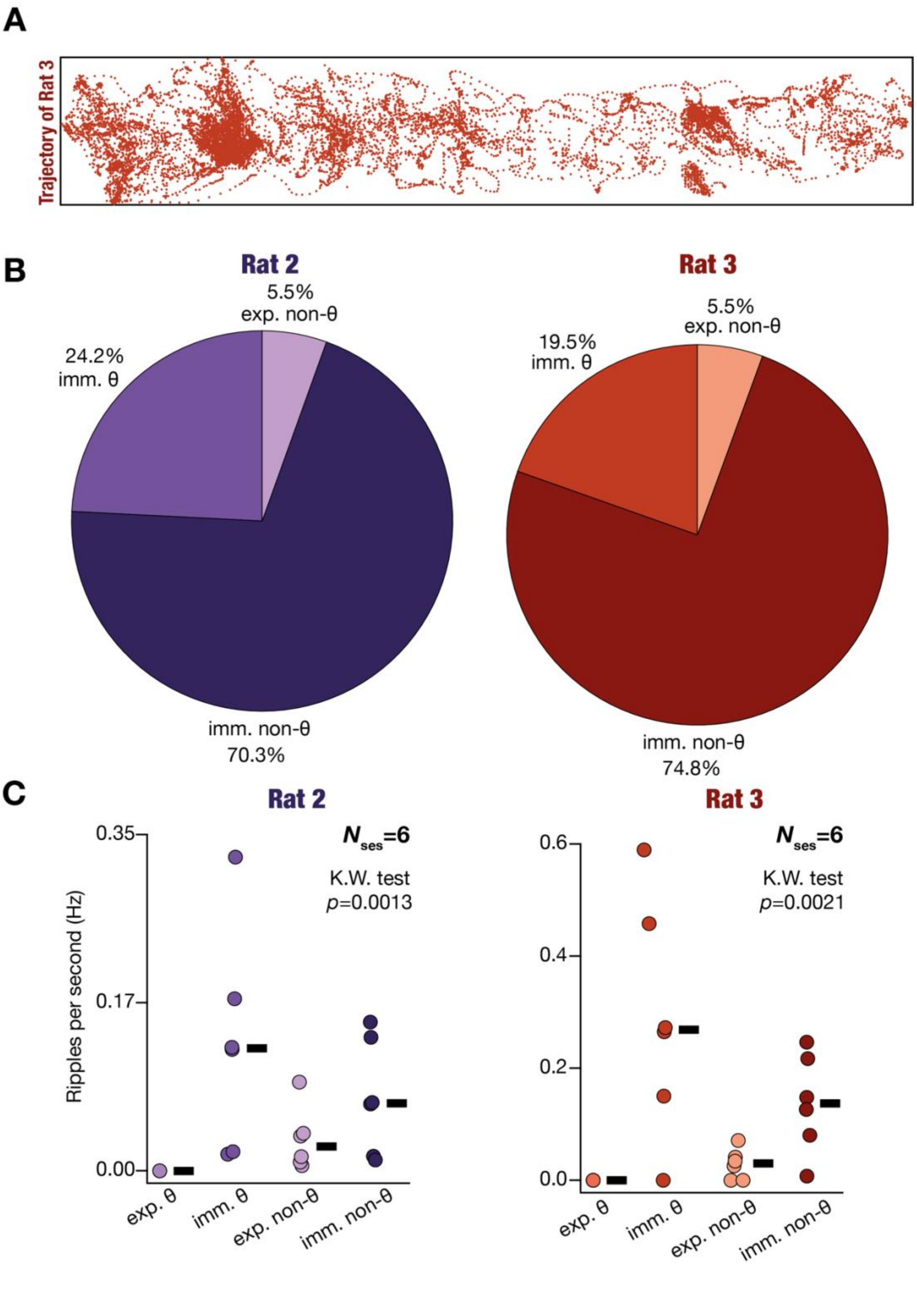

**Supplementary Figure S9. Fraction and frequency of different kinds of ripples in linear maze recordings. (A)** Linear maze trajectory of a rat in an example recording session. **(B)** Pie charts representing proportions of the four kinds of ripples — exploratory theta (exp. θ); immobile theta (imm. θ); exploratory non-theta (exp. non-θ); and immobile non-theta (imm. non-θ) — for the three rats spanning all recording sessions. **(C)** Frequency of occurrence of ripples for the four ripple kinds, provided for all recorded sessions. The thick black lines represent the respective median values across all sessions. The *p* values computed with the Kruskal Wallis (K.W.) test are provided in the figure for each rat.

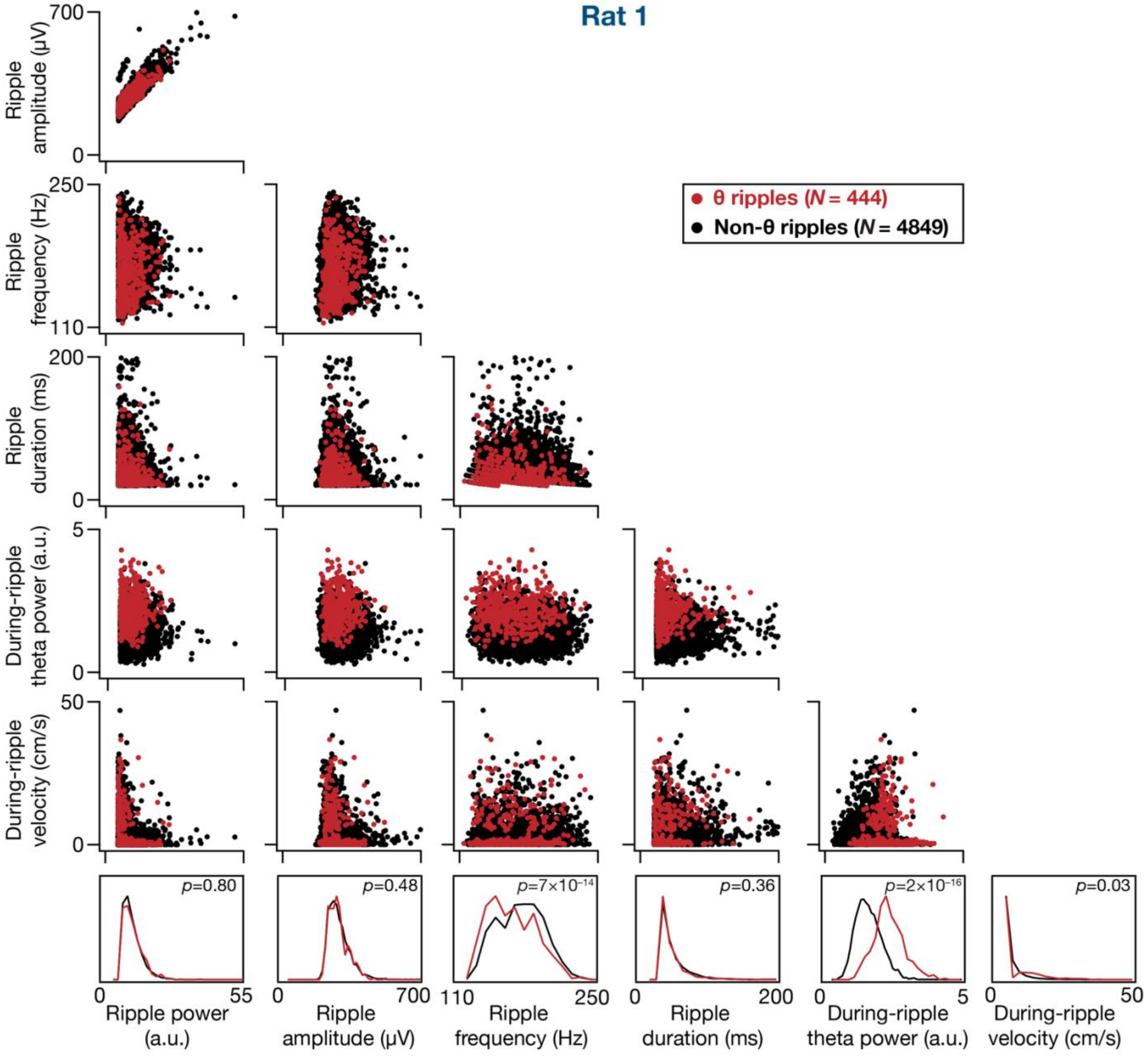

**Supplementary Figure S10. Comparing pairwise relationships between characteristics of theta and non-theta ripples for rat 1.** Scatter plots of pairwise dependencies between six characteristic measurements pertaining to ripples: power, amplitude, frequency, duration, theta power during ripple, and velocity of the rat during ripple. The theta power during ripple was taken to be the average power in the theta frequency band (6–10 Hz) in the period spanning 50 ms on either side of the peak of the ripple. Velocity during the ripple was taken to be the average velocity of the rat 150 ms on either side of the ripple peak. The last row provides the histograms of individual measurements for both theta and non-theta ripples. *p* values were computed with the Wilcox rank sum test. The Pearson correlation coefficient values were low (<0.2) for other pairwise plots except for the top left graph showing ripple amplitude *vs*. ripple power (*R* value for theta ripples: 0.89; *R* value for non-theta ripples: 0.88).

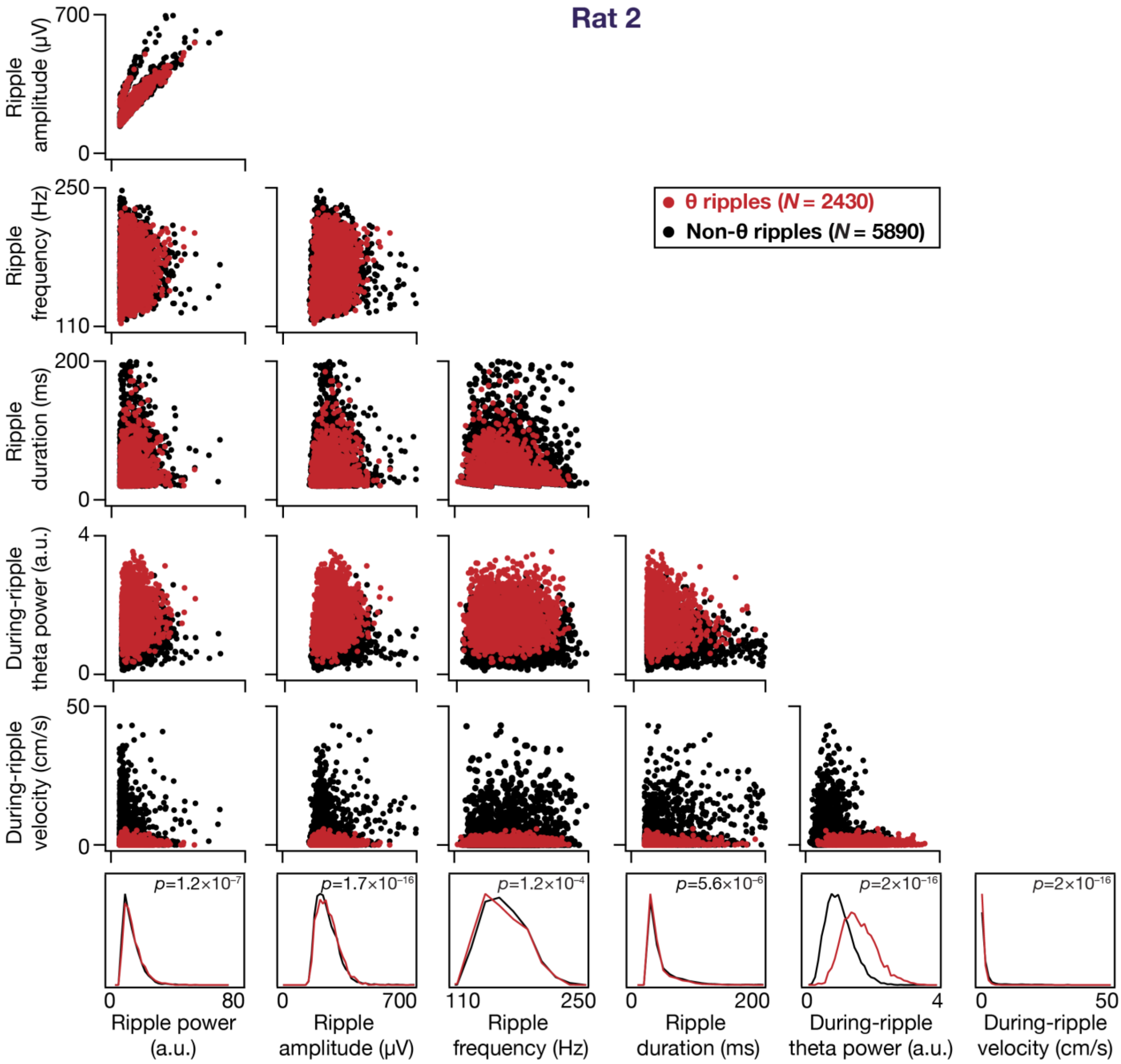

**Supplementary Figure S11. Comparing pairwise relationships between characteristics of theta and non-theta ripples for rat 2.** Scatter plots of pairwise dependencies between six characteristic measurements pertaining to ripples: power, amplitude, frequency, duration, theta power during ripple, and velocity of the rat during ripple. The theta power during ripple was taken to be the average power in the theta frequency band (6–10 Hz) in the period spanning 50 ms on either side of the peak of the ripple. Velocity during the ripple was taken to be the average velocity of the rat 150 ms on either side of the ripple peak. The last row provides the histograms of individual measurements for both theta and non-theta ripples. *p* values were computed with the Wilcox rank sum test. The Pearson correlation coefficient values were low (<0.2) for other pairwise plots except for the top left graph showing ripple amplitude *vs*. ripple power (*R* value for theta ripples: 0.89; *R* value for non-theta ripples: 0.87).

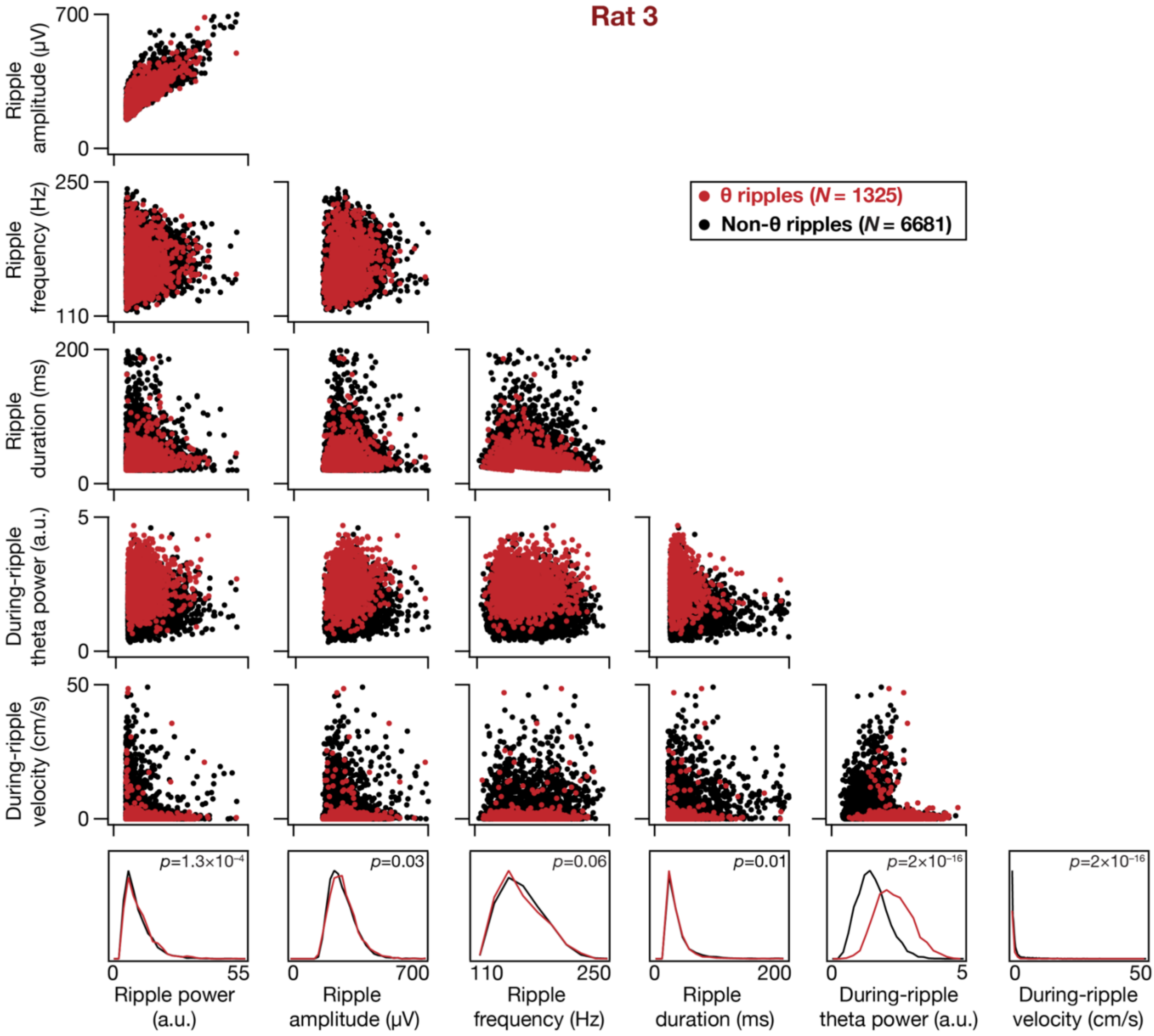

**Supplementary Figure S12. Comparing pairwise relationships between characteristics of theta and non-theta ripples for rat 3.** Scatter plots of pairwise dependencies between six characteristic measurements pertaining to ripples: power, amplitude, frequency, duration, theta power during ripple, and velocity of the rat during ripple. The theta power during ripple was taken to be the average power in the theta frequency band (6–10 Hz) in the period spanning 50 ms on either side of the peak of the ripple. Velocity during the ripple was taken to be the average velocity of the rat 150 ms on either side of the ripple peak. The last row provides the histograms of individual measurements for both theta and non-theta ripples. *p* values were computed with the Wilcox rank sum test. The Pearson correlation coefficient values were low (<0.2) for other pairwise plots except for the top left graph showing ripple amplitude *vs*. ripple power (*R* value for theta ripples: 0.82; *R* value for non-theta ripples: 0.83).

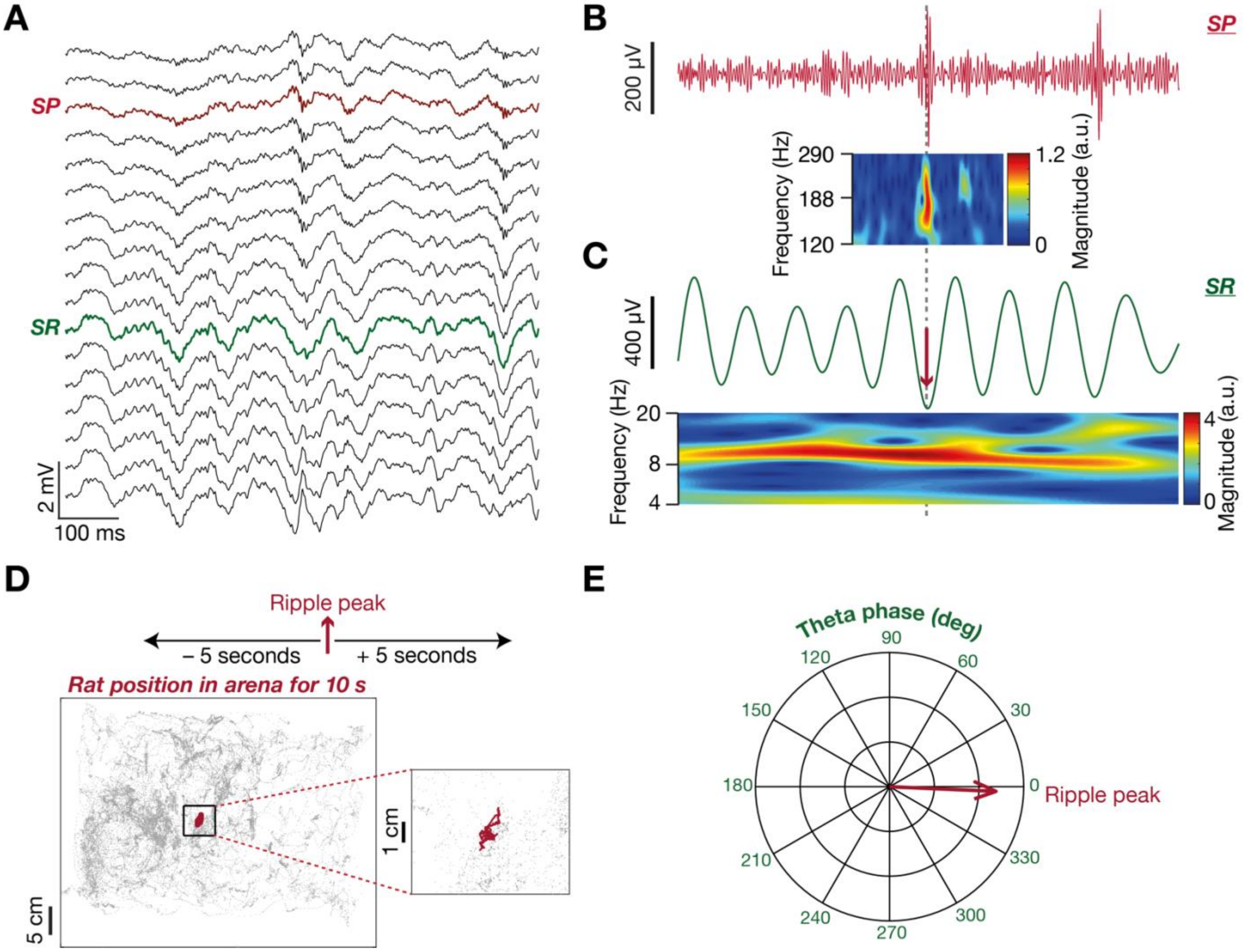

**Supplementary Figure S13. An example immobile theta (imm. θ) ripple. (A)** Laminar profile of local field potentials recorded during an example session from rat 1 with the identified SP and SR marked in red and green respectively. **(B)** Ripple-filtered trace and ripple-frequency wavelet scalogram of the signal recorded in SP. **(c)** Theta-filtered trace and theta-frequency wavelet scalogram of the signal recorded in SR. Simultaneous prevalence of ripple in SP and theta in SR may be noted both from the laminar profile (A) and *strata*-wise scalograms (B–C). **(D)** Trajectory of the rat 5 seconds before and after the ripple peak time shown in red overlaid on top of the trajectory of the rat during the entire session in grey. The zoomed inset shows the position of the rat during the 10 s period. **(E)** Phase of the peak of the ripple shown in (B) detected in SP with respect to the trough of the simultaneously recorded theta oscillation shown in (C) detected in SR.

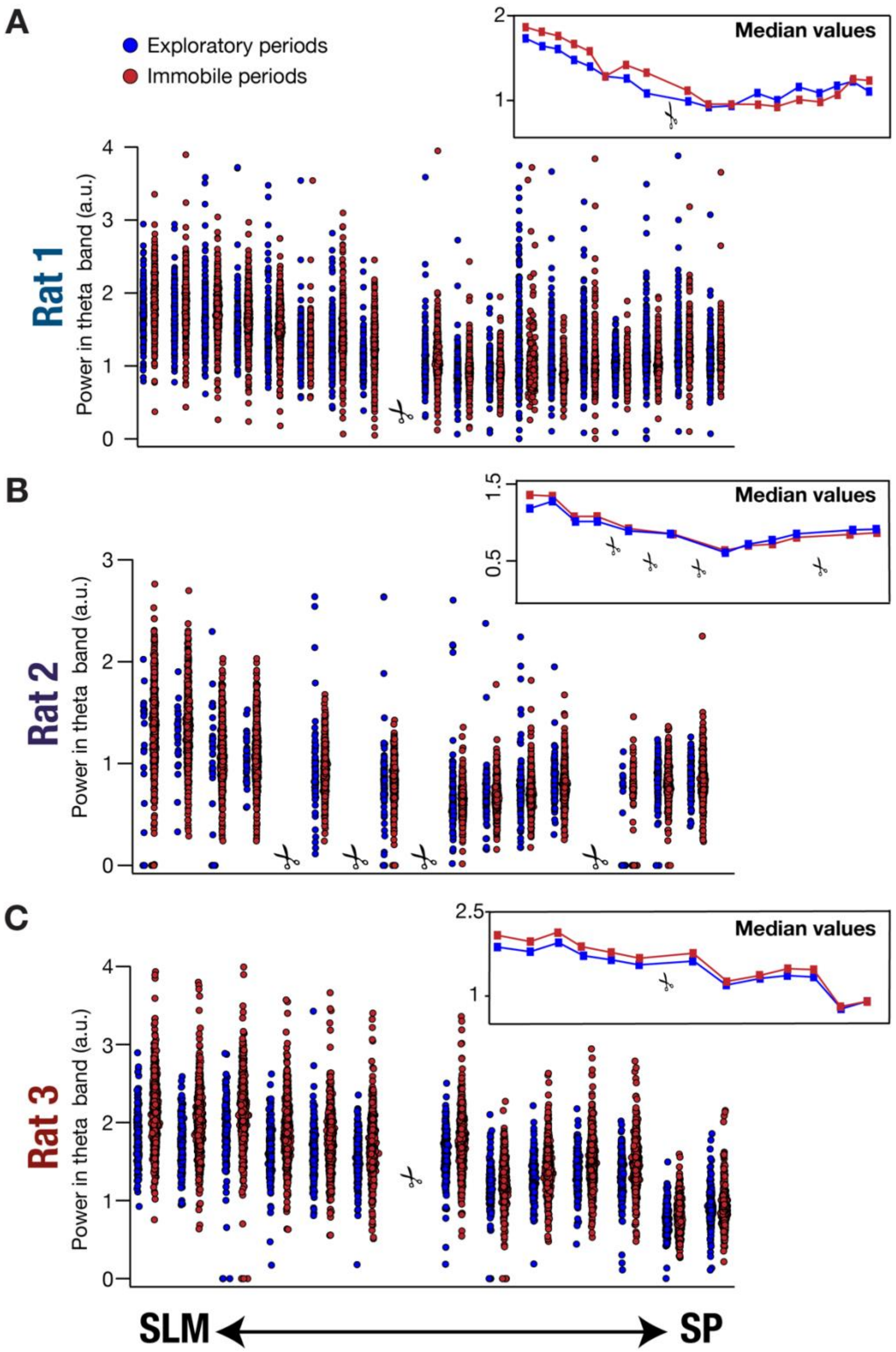

**Supplementary Figure S14. Theta power was highest in SLM with a progressive power reduction through SR to SP during immobile and exploratory periods. (A–C)** Theta power during immobile and exploratory periods obtained from all detected theta epochs from recording channels spanning different *strata* (SLM to SP) for the three rats across all recording sessions. The insets to the right show the *stratum*-dependent median power values computed from the respective points shown in the main plots for both immobile and exploratory periods. A symbol showing a pair of scissors indicates that the specific recording channel was noisy for a majority (≥ 90%) of the recorded sessions and was discarded from analyses. Wilcox rank sum test *p* values for theta power computed from the identified SLM channel *vs*. SP channel: **Rat 1**: $2.2\times10^{-16}$ (exp.), $2.2\times10^{-16}$ (imm.); **Rat 2**: $3.9\times10^{-11}$ (exp.), $2.2\times10^{-16}$ (imm.); **Rat 3**: $2.2\times10^{-16}$ (exp.), $2.2\times10^{-16}$ (imm.). These panels expand on the summary data provided in Fig. 6. Whereas data points in Fig. 6 represent average values of theta power in each session, data points here correspond to theta power in single theta epochs.

**Supplementary Table S1: Controls for computing theta ripple co-occurrences.** The crucial parameters that were used in identifying theta ripples and behavioral state were varied to assess the implications of changing these parameters. Reported are the variations in the numbers of theta ripples, exploratory theta ripples and immobile theta ripples for parametric combination. The parameters varied were as follows:

1. **Maze type** – All the experiments were performed in the open box arena; a linear maze was used as a control for evaluating our hypothesis in a different arena.
2. **Theta frequency range** – All the analyses were done with a theta frequency range of 6–10 Hz. A lower range of 4 Hz (instead of 6 Hz) was also used as a control.
3. **Velocity thresholds** – A Default value of 5 cm/sec was used as both the minimum and maximum velocity threshold for consideration as an exploratory and immobile period respectively. Three other velocity thresholds (4cm/sec; 6cm/sec and 8cm/sec) were used as controls.
4. **Ripple Validation** – A default value of 10 was used as the number of channels in which ripple-like events were detected and compared in order to discard artifacts from the *stratum pyramidale.* Specifically, putative ripple events in the SP layer were discarded if there were events detected in at least 10 channels within 10 ms on either side of the ripple under consideration. As a control for this number, 12 channels were used for comparing and discarding artifacts from SP.
5. **Theta detection** – To quantify the strength of theta detection, a default of 1 SD above the mean signal amplitude was used as the theta detection threshold (**medium** level). As a control, 0.5 SD and 1.5 SD above the mean signal amplitude were also used as controls to signify **low** and **high** levels of detection thresholds.

Default parametric combination used for analyses (listed as “Default” below):

> [<u>*Maze type*</u>: **Open Box**; <u>*Theta frequency range*</u>: **[6 10] Hz**; <u>*Velocity thresholds*</u>: [**5 cm/s** (exp. Min. velocity), **5 cm/s** (Imm. Max. velocity); <u>*Ripple Validation*</u>: **10 channels**; <u>*Theta detection*</u>: **Medium** (1 SD above mean signal amplitude) **level**]

| Parameter changed | # Theta ripples | # Exploratory theta ripples | # Immobile theta ripples |
|---|---|---|---|
| **RAT #1** | | | |
| **Default** | 444 | 24 | 312 |
| **Maze type (Linear Maze)** | – | – | – |
| **Theta frequency range ([4 10] Hz)** | 607 | 18 | 498 |
| **Velocity Threshold (4,4)** | 444 | 39 | 303 |
| **Velocity Threshold (6,6)** | 444 | 18 | 313 |
| **Velocity Threshold (8,8)** | 444 | 6 | 316 |
| **Ripple Validation (# Channels: 12)** | 444 | 39 | 303 |
| **Theta detection (Low level)** | 915 | 31 | 664 |
| **Theta detection (Medium level)** | 204 | 14 | 145 |
| **RAT #2** | | | |
| **Default** | 2430 | 0 | 2261 |
| **Maze type (Linear Maze)** | 68 | 0 | 53 |
| **Theta frequency range ([4 10] Hz)** | 2568 | 1 | 2373 |
| **Velocity Threshold (4,4)** | 2430 | 0 | 2185 |
| **Velocity Threshold (6,6)** | 2430 | 0 | 2347 |
| **Velocity Threshold (8,8)** | 2430 | 0 | 2410 |
| **Ripple Validation (# Channels: 12)** | 2586 | 1 | 2405 |
| **Theta detection (Low level)** | 3856 | 3 | 3567 |
| **Theta detection (Medium level)** | 1405 | 0 | 1314 |
| **RAT #3** | | | |
| **Default** | 1325 | 11 | 1206 |
| **Maze type (Linear Maze)** | 110 | 0 | 74 |
| **Theta frequency range ([4 10] Hz)** | 1615 | 1 | 1488 |
| **Velocity Threshold (4,4)** | 1325 | 14 | 1167 |
| **Velocity Threshold (6,6)** | 1325 | 6 | 1259 |
| **Velocity Threshold (8,8)** | 1325 | 1 | 1283 |
| **Ripple Validation (# Channels: 12)** | 1357 | 14 | 1233 |
| **Theta detection (Low level)** | 2402 | 23 | 2130 |
| **Theta detection (Medium level)** | 701 | 3 | 649 |
